# ER stress-induced TREM2 downregulation exacerbates platelet activation and myocardial infarction in patients with coronary artery disease

**DOI:** 10.1101/2024.08.20.608887

**Authors:** Xiaowen Wu, Guanxing Pan, Lin Chang, Yulong Zhang, Yangyang Liu, Wei Zhang, Yifan Guo, Ge Zhang, Haoxuan Zhong, Zhiyong Qi, Jianjun Zhang, Ruyi Xue, She Chen, Hu Hu, Jianzeng Dong, Si Zhang, Zhongren Ding

## Abstract

Coronary artery disease (CAD) is characterized by the chronic immune-inflammation, excessive endoplasmic reticulum (ER) stress, and platelet hyperactivity; however, whether there is a signaling hub linking these events remains unclear. Here, we identified that triggering receptor expressed on myeloid cells 2 (TREM2), an important pattern recognition receptor of the innate immune system, may serve as one such hub. We found that platelets expressed TREM2 and platelets from CAD patients had decreased TREM2 expression compared to healthy subjects. Decreased TREM2 is associated with platelet hyperactivity in CAD patients. This decrease could be due to excessive ER stress, which downregulated TREM2 through the CHOP-C/EBPα axis. Loss of TREM2 not only enhanced platelet activation in response to ADP, collagen, and collagen-related peptide (CRP), but also amplified the platelet inflammatory response. Loss of TREM2 exacerbated mouse mesenteric arterial thrombosis and aggravated experimental myocardial infarction (MI). Moreover, a TREM2-activating antibody inhibited platelet activation, alleviated arterial thrombosis and pulmonary embolism. In addition, TREM2-activating antibody exhibited cardioprotective roles against experimental MI and reduced the inflammatory burden. Mechanistically, TREM2/DAP12/SHIP1 axis negatively regulated platelet activation through reducing PIP3 levels and inhibiting Akt phosphorylation. We also provided evidence supporting sphingosine-1-phospage (S1P) as a physiological agonist of TREM2. In summary, we find that TREM2 connects chronic immune-inflammation, excessive ER stress, and platelet hyperactivity in CAD patients. Downregulating TREM2 by ER stress exacerbates platelet activation and amplifies inflammation response in patients with CAD. TREM2-activating antibodies may have therapeutic potential for CAD patients.

**Key Points:**

- Platelets from CAD patients have decreased TREM2 expression, which is caused by ER stress and associated with platelet hyperactivity.
- S1P-TREM2-SHIP1 pathway inhibits platelet activation, alleviates arterial thrombosis, and exhibits cardioprotective roles against experimental myocardial infarction.

Visual Abstract.
Downregulation of platelet TREM2 caused by ER stress in CAD patients leads to platelet hyperactivity and aggravates myocardial ischemia, which is rescued by TREM2-activating antibody.
Excessive ER stress in CAD upregulates CHOP, which dimerizes with C/EBPα, a transcription factor of TREM2, decreases TREM2 transcription and expression in megakaryocytes and further in platelets. TREM2/DAP12/SHIP1/Akt pathway negatively regulates platelet activation; decreased TREM2 thus aggravates platelet hyperactivity, inflammation, and myocardial infarction, which can be rescued by TREM2 activation including antibody or the small molecule agonist. Physiologically, S1P released from platelet α granules is an endogenous agonist of TREM2.

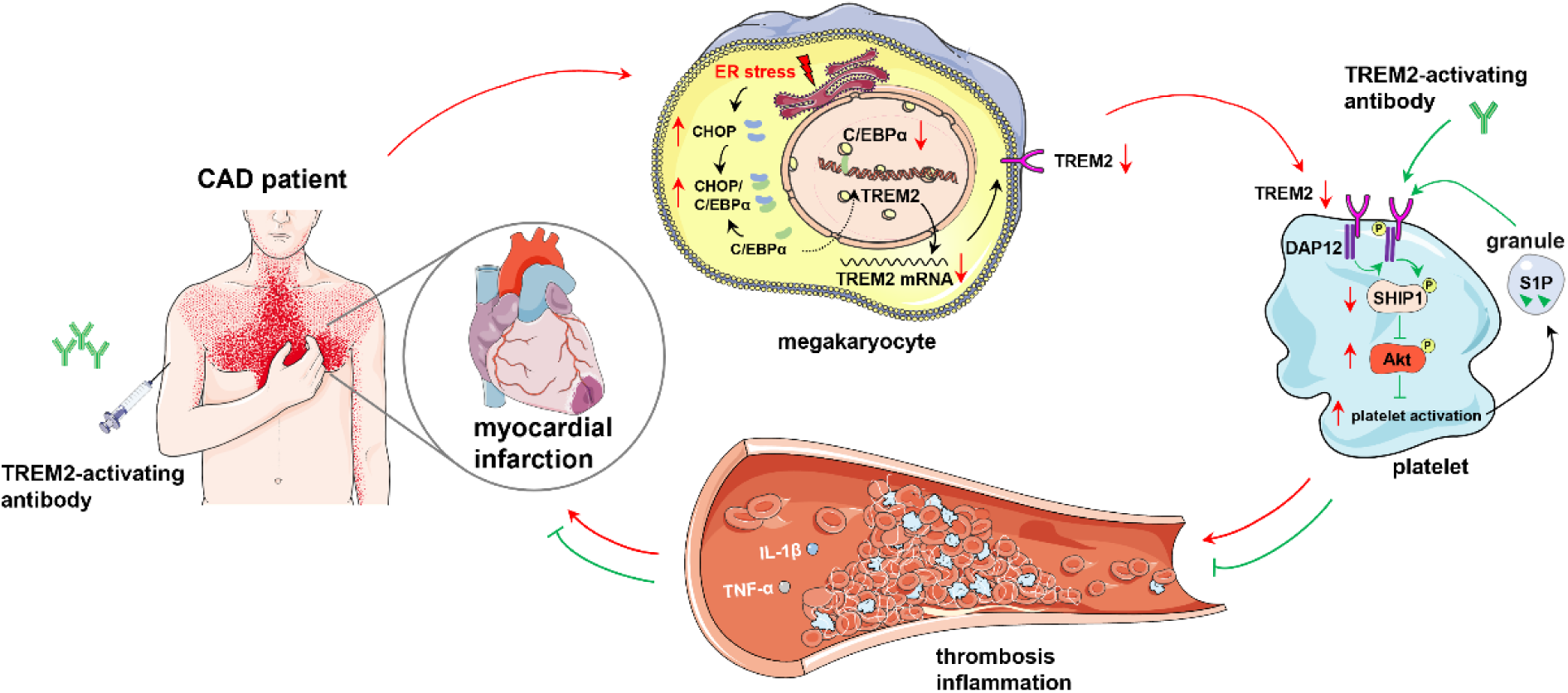

## Introduction

Coronary artery disease (CAD), the most common atherothrombotic disease, is the leading cause of death worldwide^1^. CAD is characterized by the platelet hyperactivity, excessive endoplasmic reticulum (ER) stress and chronic immune-inflammation. Platelet hyperactivity plays a critical role in CAD in both pathogenesis of atherosclerosis and development of acute thrombotic events. Increased platelet activation promotes the initiation and progression of atherosclerosis, and consequently induces occlusive thrombus formation in the coronary arteries, ultimately leading to acute coronary syndrome (ACS)^2,3^. Recently, there is growing evidence that various conditions associated with CAD could induce excessive cellular endoplasmic reticulum (ER) stress. Meanwhile, ER stress further evokes the onset and aggravates the development of CAD, creating a rather vicious cycle^4,5^. In addition to ER stress, CAD is coupled with a chronic inflammatory process driven by the innate immune response. The innate immune response is initiated by pattern recognition receptors (PRRs). PRRs recognize evolutionarily conserved pathogen-associated molecular patterns (PAMPs) and damage-associated molecular patterns (DAMPs). Pathogen-released PAMPs can initiate immune inflammatory responses in atherosclerosis, and the subsequent tissue damage will generate many DAMPs to further influence the progression of atherosclerosis. We and others have reported that PRRs including toll-like receptors (TLRs) and nucleotide-binding oligomerization domain-like receptors (NLRs) are expressed in platelets and act as a bridge linking innate immunity and platelet activation^6–8^. Stimulation of these PRRs generally enhances platelet activation and thrombosis; however, the PRRs responsible for the negative regulation of platelet activation remain unknown. In addition, it is unclear whether there is a link between ER stress, innate immunity, and platelet hyperactivity in CAD.

In this study, we found that platelets express TREM2, an important PRR previously reported in myeloid cells such as macrophages and monocyte-derived dendritic cells^9^, osteoclasts^10^, and microglia^11^. In recent years, TREM2 has been established as an immune signaling hub that senses tissue damage in metabolic syndrome, Alzheimer’s disease, and cancer^12,13^. Here, we identified TREM2 as a signaling hub linking innate immunity, ER stress, and platelet hyperactivity in CAD patients. Excessive ER stress in CAD patients downregulates TREM2 expression through the CHOP-C/EBPα axis in platelets. In turn, downregulated TREM2 increases platelet activation, thrombosis, inflammation, and experimental MI through the DAP12/SHIP1/Akt axis. We also proposed that S1P released from platelet α granules as a physiological agonist of TREM2. Importantly, we found that the TREM2 activating antibody (a treatment strategy tested in clinical trial for Alzheimer’s disease, NCT04592874), which exhibits antiplatelet, anti-inflammatory, antithrombotic, and cardioprotective roles against experimental MI, may serve as a new therapeutic agent for atherosclerotic diseases.

## Methods

Detailed Materials and Methods are described in the Supplemental Material Online.

### Mice

TREM2 deficient mice on C57BL/6J background were obtained from the Jackson Laboratory (Bar Harbor, ME). Mice aged 8 - 10 weeks were used. All animal procedures were performed according to the criteria outlined in the “Guide for the Care and Use of Laboratory Animals” prepared by the National Academy of Sciences and published by the National Institutes of Health (NIH publication 86 - 23 revised 1985).

### Human subjects

All experiments using human blood samples were performed in accordance with the Declaration of Helsinki and approved by the Institutional Review Board of Fudan University.

### Data presentation

Data were expressed as the mean ±SEM, and median with 25 - 75 percentiles was used for data with no-Gaussian distribution. Differences were considered significant at *P* < 0.05. Details on statistical analysis were presented in the Supplementary Material Online.

## Results

### Platelets express TREM2 and platelet TREM2 is significantly downregulated in CAD patients

We found that human and murine platelets robustly express TREM2 at both mRNA and protein levels, as detected by RT-PCR (Figure 1A) and Western blot (Figure 1B). Immunofluorescence staining further demonstrated that TREM2 was localized to both the membrane and cytoplasm of platelets (Figure 1C). Platelets from patients with CAD, including SAP and ACS, expressed significantly lower levels of TREM2 at both the protein and mRNA levels (Figure 1D and E) than those from healthy donors. Moreover, ACS patients had the lowest TREM2 expression (Figure 1D and E), suggesting that decreased platelet TREM2 expression is associated with the severity of CAD. The impaired expression of TREM2 in platelets from patients with CAD was recapitulated by flow cytometry (Figure 1Fi). Using platelets from 25 patients with ACS, we further found that the expression of TREM2 was negatively correlated with CRP-induced P-selectin (CD62P) release from platelet α granules (Figure 1Fii). Together, these data suggest that decreased TREM2 expression is associated with platelet hyperactivation in CAD patients.

**Figure 1.**
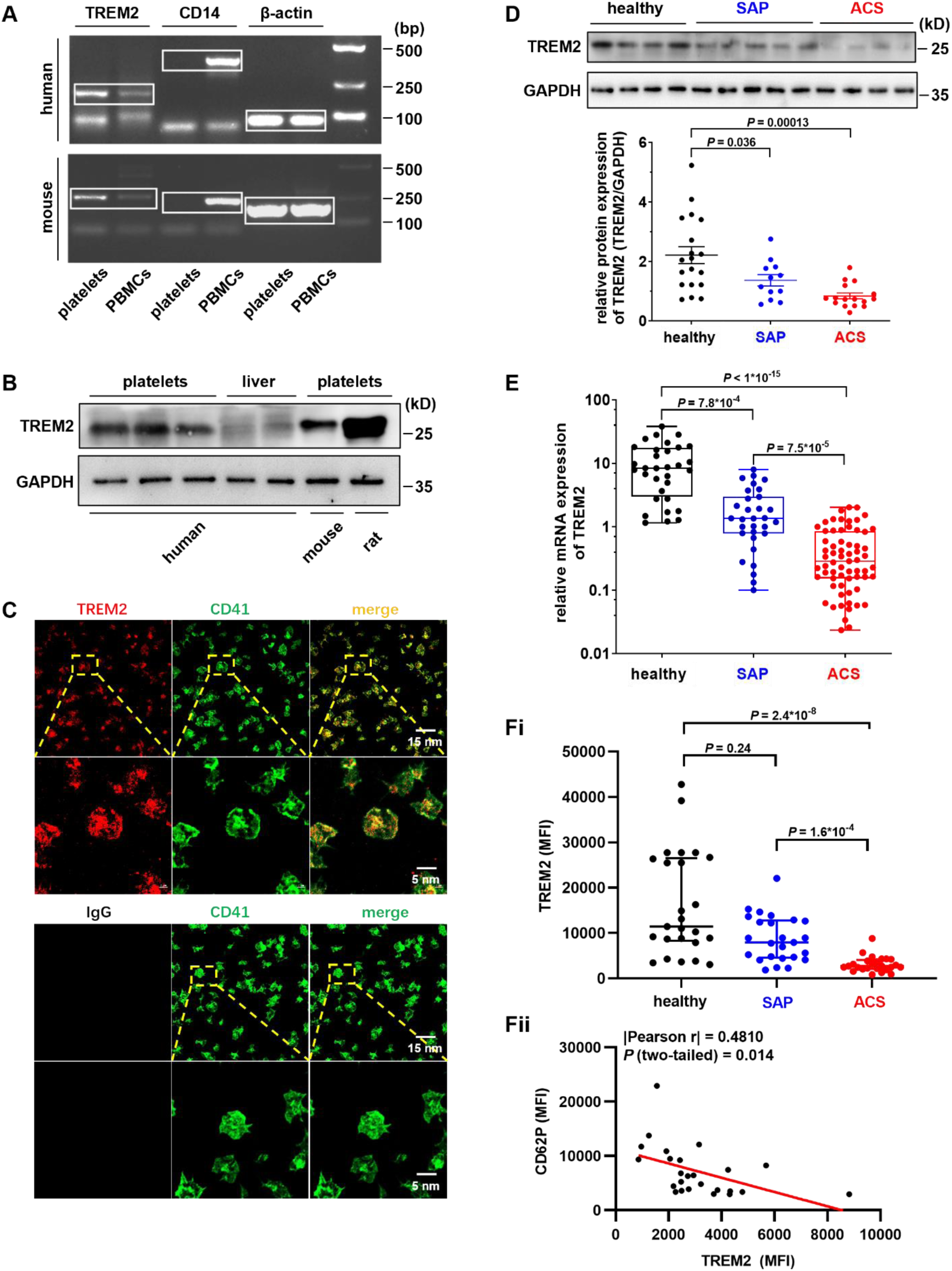
Platelets express TREM2, which is decreased in patients with CAD. **A.** RT-PCR detection of TREM2 mRNA in platelets and peripheral blood mononuclear cells (PBMCs). Monocyte-specific marker CD14 was used to rule out contamination of white blood cells. Human 227 bp of TREM2, 357 bp of CD14, and 90 bp of β-actin are amplified; mouse 260 bp of TREM2, 247 bp of CD14, and 174 bp of β-actin are amplified. **B.** Platelets express TREM2 protein. Human liver tissue (non-tumor liver tissue) from a patient with hepatocellular carcinoma was used as a negative control. **C.** Expression of TREM2 in human platelets as detected by confocal microscopy. Platelets were stained with antibodies against TREM2 (IgG as isotype control) and platelet marker CD41, and then detected using Alexa 594 and Alexa 488-labeled secondary antibodies, respectively. **D.** Decreased TREM2 protein levels in platelets from patients with CAD. The typical Western blot result and statistical analysis of 19 healthy donors, 12 patients with SAP, and 16 patients with ACS are shown. **E.** Decreased TREM2 mRNA expression in platelets from patients with SAP (n = 31) and ACS (n = 65) compared with healthy donors (n = 32), as evaluated by qPCR. mRNA levels were normalized to β-actin. **F.** Platelets from patients with ACS express lower TREM2, which is reversely correlated with CRP-induced P-selectin release analyzed by flow cytometry. **i)**. Decreased platelet TREM2 expression in patients with ACS compared with the healthy and patients with SAP analyzed by flow cytometry (n = 25 in each group). **ii)**. CRP-induced platelet P-selectin (CD62P) release in patients with ACS is reversely correlated with platelet TREM2 expression level. Platelets from the same 25 patients with ACS in panel **Fi** were stimulated by 0.1 µg/mL CRP, P-selectin release was analyzed by flow cytometry. Each spot represents a different individual. Characteristics of the study population D, E, and F are summarized in Supplement Table III, IV, and V respectively. One-way ANOVA followed by Tukey’s multiple comparison test was performed in (**D**). Kruskal-Wallis test followed by Dunn’s multiple comparison test was performed in **(E)** and **(Fi)**. Pearson’s correlation analysis was used in (**Fii**).

### Elevated transcriptional repressor CHOP inhibits C/EBPα-mediated TREM2 transcription in CAD patients

Genetic alteration and epigenetic inactivation are two major causes of protein downregulation. Therefore, we went on to detect the sequence of the TREM2 exons in 50 ACS patients (Supplemental Table VI) using Sanger sequencing. We found that neither the mutations nor methylation status is the primary mechanism regulating TREM2 downregulation (Supplemental Figure 1A and B).

Next, we investigated whether transcriptional regulation was responsible for the downregulation of TREM2 in platelets from CAD patients. To define the minimal DNA sequence required for TREM2 promoter activity, we evaluated the transcriptional activity of a series of TREM2 promoter truncation constructs transfected into HEK293T cells using a luciferase reporter assay. We found that the upstream sequence, −400 to −201, is the minimal sequence with the highest relative luciferase activity (Supplemental Figure 2A). Next, using P-Match (http://www.gene-regulation.com/cgi-bin/pub/programs/pmatch/bin/p-match.cgi), a program predicting transcription factor binding sites, we found that the −400 to −201 region contains a consensus binding site, ^-298^TTGCA^-294^, for the transcription factor CCAAT enhancer-binding protein α (C/EBPα, Figure 2A).

**Figure 2.**
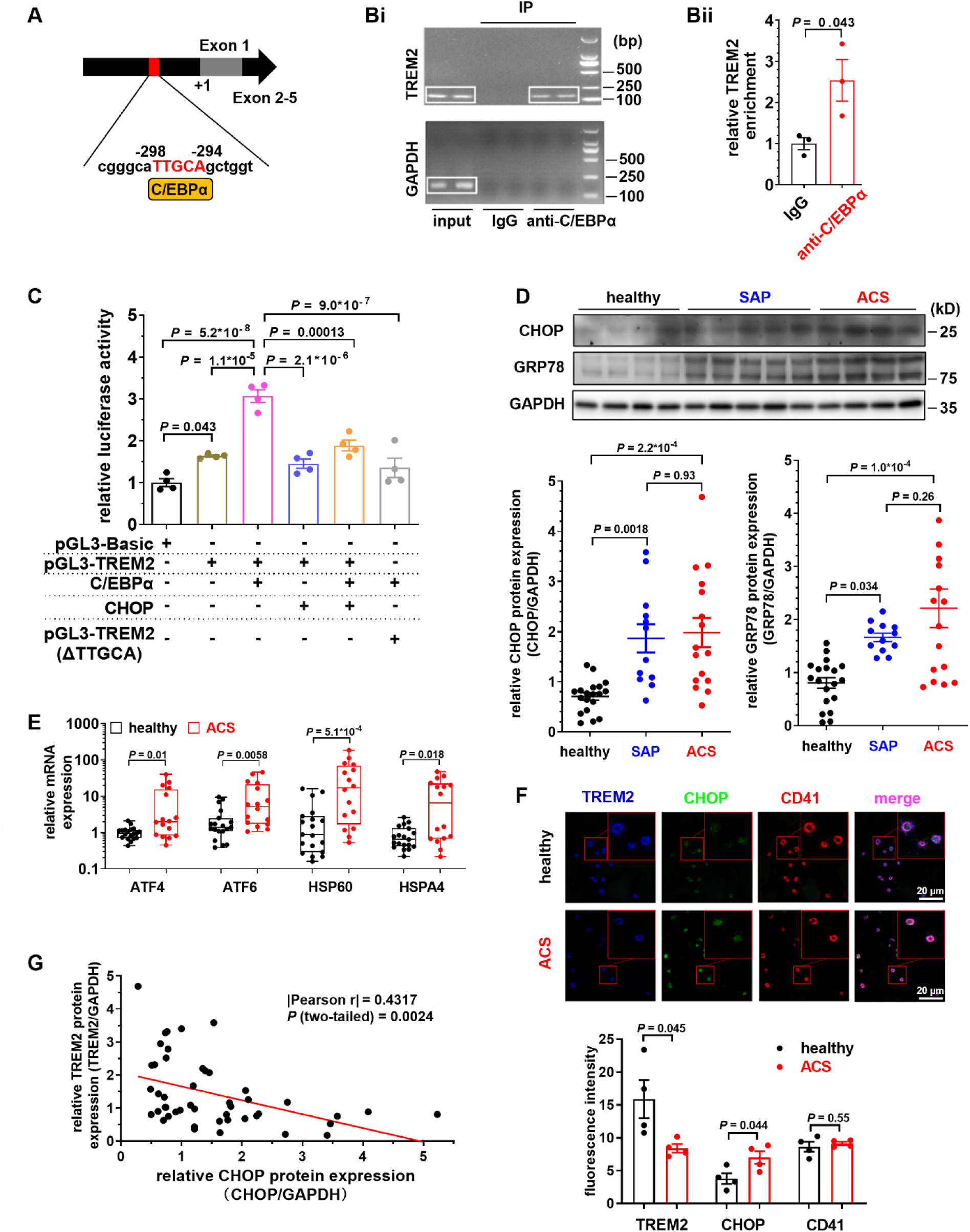
Increased CHOP inhibits C/EBPα-mediated TREM2 transcription in CAD. **A.** Predicted C/EBPα binding site in human TREM2 promoter. **B**. ChIP analysis of C/EBPα binding to the TREM2 promoter in Meg-01 cells. **i)**. PCR amplification of TREM2 promoter region containing C/EBPα binding motif from Meg-01 cells after ChIP. Nonspecific lgG was used as a control. **ii)**. qPCR analysis of C/EBPα-bound TREM2 promoter from Meg-01 cells after ChIP. Data are normalized to the pre-immunoprecipitation input for each sample and expressed as the fold change (n = 3). **C.** Luciferase reporter assay in 293T cells revealed that C/EBPα binds to the ^-298^TTGCA^-294^ region of the TREM2 promoter to promote transcription, which is inhibited by CHOP and deletion of the ^-298^TTGCA^-294^ region (^Δ^TTGCA) (n = 4). **D.** Increased CHOP and GRP78 expression in platelets from patients with SAP and ACS. Representative Western blot result and a summary of the data from 19 healthy donors, 12 SAP and 16 ACS patients are provided. **E.** Increased mRNA expression of ATF4/6 and HSP60/A4 in platelets from patients with ACS as detected by qPCR. The mRNA level was normalized to β-actin. Data from 19 healthy donors and 16 ACS patients were presented. **F.** Decreased TREM2 was accompanied by increased CHOP in platelets from patients with ACS. Representative immunofluorescence images and the summary (n = 4) are shown. **G.** CHOP expression is negatively correlated with TREM2 protein expression in platelets. Samples from Figures 1D & 2D including 19 healthy subjects, 12 SAP, and 16 ACS patients were analyzed (n = 47). Characteristics of the study population are provided in Supplemental Table III. Unpaired t-test was performed in (**B**) and (**F**). Statistical analyses were performed using one-way ANOVA followed by Tukey’s multiple comparison test in **(C)** and **(D)**. Mann-Whitney test was performed in (**E**). Pearson’s correlation was used in (**G**).

We then used Meg-01, a human megakaryoblastic cell line, to investigate the transcriptional regulation of TREM2. Chromatin immunoprecipitation (ChIP) assay demonstrated that C/EBPα was enriched at the TREM2 promoter in Meg-01 cells (Figure 2Bi). qPCR-based CHIP assay showed that C/EBPα was enriched in TREM2 promoter by 2.5 folds compared with IgG (Figure 2Bii). Luciferase reporter assay further confirmed that C/EBPα increased the luciferase reporter activity of TREM2 promoter by 1.87 folds. The ^-298^TTGCA^-294^ region is necessary for the binding of C/EBPα to TREM2 promoter because the deletion of the ^-298^TTGCA^-294^ region (^Δ^TTGCA) abolished C/EBPα-induced TREM2 promoter activity (Figure 2C).

C/EBP-homologous protein (CHOP) is a classic biomarker and mediator of the ER stress response^14–16^. It dimerizes with C/EBPα and inhibits the binding of C/EBPα to its promoter-binding site^17^. Consistently, we found that CHOP inhibited C/EBPα-triggered TREM2 promoter luciferase reporter activity (Figure 2C). Moreover, the electrophoretic mobility shift assay (EMSA) demonstrated that C/EBPα directly binds to the C/EBPα-response element (^-298^TTGCA^- 294^) in TREM2 promoter, which is prevented by the addition of purified CHOP protein (Supplemental Figure 2B). We also observed a significant increase in CHOP protein, which inhibits TREM2 transcription, in CAD patient platelets (Figure 2D). Taken together, these observations indicate that elevated CHOP may inhibit TREM2 gene transcription by preventing C/EBPα from binding to TREM2 promoter in CAD patients.

### Increased platelet ER stress in CAD patients

Increased CHOP expression indicated severe ER stress, which plays a pivotal role in the development of cardiovascular disease, including atherosclerosis, ischemic heart disease, and heart failure^14^. Previous studies have reported that monocytes, cardiomyocytes, and endothelial cells from CAD patients undergo increased ER stress^18,19^. However, it is unclear whether excessive ER stress is present in the platelets from patients with CAD. We found that the 78 kDa glucose-regulated protein (GRP78), another marker of ER stress^14^, was increased in platelets from CAD patients, evidenced by western blot (Figure 2D). In addition, qPCR assays revealed a dramatic increase in ER stress sensors, including activating transcription factor 4/6 (ATF4/6), heat shock protein 60 (HSP60), and heat shock protein family A (Hsp70) member 4 (HSPA4) in platelets from ACS patients (Figure 2E). Immunofluorescence analysis further confirmed the increased CHOP and decreased TREM2 expression in platelets from ACS patients compared to platelets from healthy subjects (Figure 2F). Finally, Pearson correlation analysis revealed a negative correlation between CHOP and TREM2 protein expression in platelets from healthy subjects and CAD patients (Figure 2G). Together, these results indicate the presence of excessive ER stress in platelets from patients with CAD.

### ER stress downregulates TREM2 via CHOP in Meg-01 cells and platelets

Next, we explored whether ER stress downregulated TREM2 expression through CHOP. Tunicamycin is a widely used inducer of ER stress. We used tunicamycin to induce ER stress in Meg-01 cells and found that the level of TREM2 decreased and the levels of CHOP and GRP78 increased, while the levels of C/EBPα remained unchanged (Figure 3A, Supplemental Figure 3A). Flow cytometry and fluorescence microscopy further confirmed that tunicamycin decreased TREM2 expression (Figure 3B and C). Moreover, tunicamycin-induced reduction in TREM2 was rescued by lentiviral shRNA-mediated silencing of CHOP in Meg-01 cells (Figure 3D and E, Supplemental Figure 3B), further confirming that ER stress downregulates TREM2 via CHOP. Consistent with the *in vitro* findings in Meg-01 cells, intraperitoneal injection of tunicamycin into mice also decreased TREM2 expression, increased CHOP and GRP78 expression, and had no effect on C/EBPα expression in megakaryocytes *in vivo* (Figure 3F).

**Figure 3.**
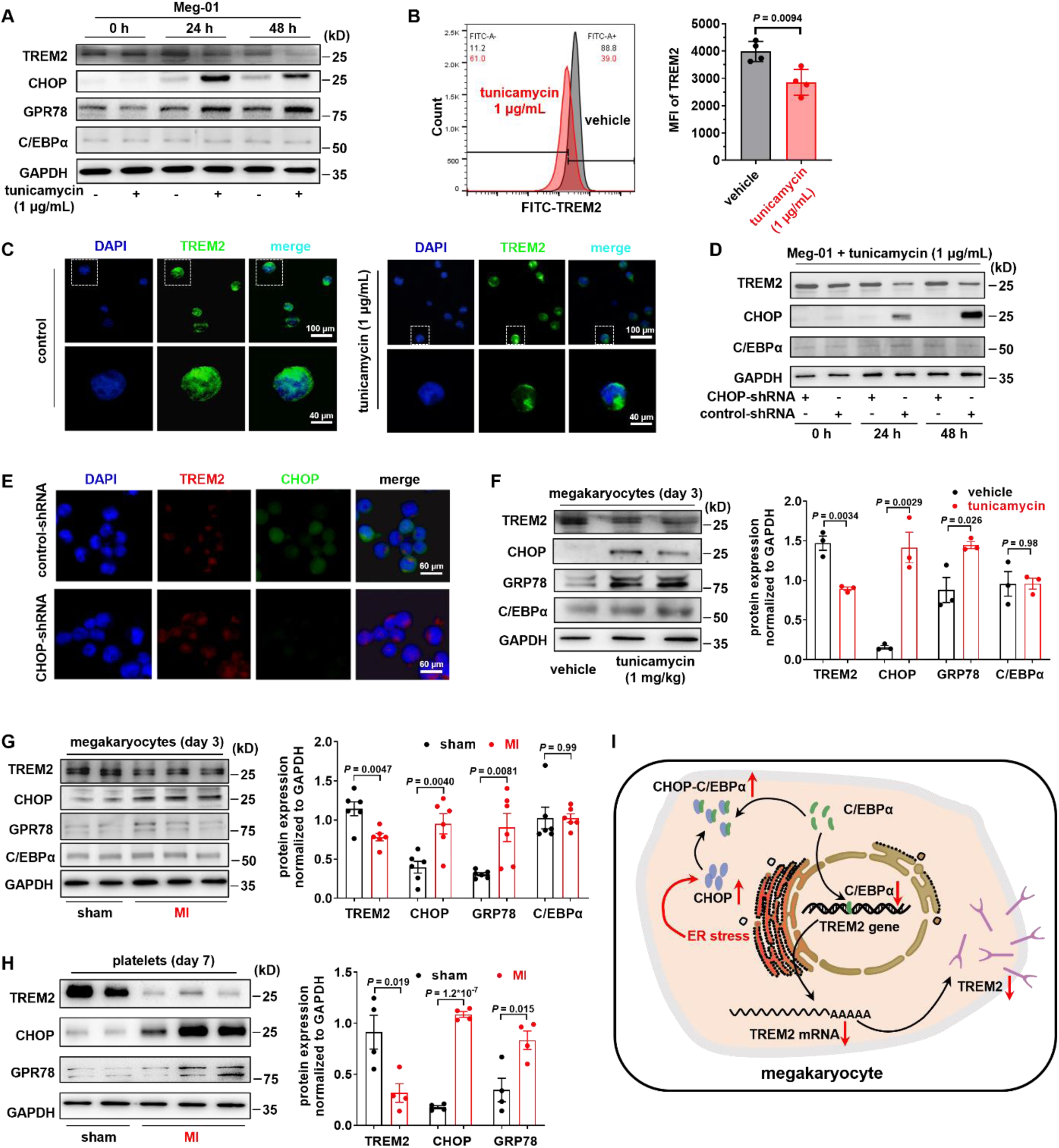
ER stress downregulates TREM2 expression in Meg-01 and platelets. **A.** Tunicamycin upregulates GRP78 and CHOP, and downregulates TREM2 expression without affecting C/EBPα levels in Meg-01 cells. Typical blots and their relative densities from 5 experiments are shown (Supplemental Figure 3A). Unless otherwise indicated, Meg-01 cells were treated with tunicamycin 1 μg/mL for 48 hours to induce ER stress. **B & C.** Tunicamycin downregulates TREM2 expression in Meg-01 cells. TREM2 expression was analyzed by flow cytometry (panel B) and immunofluorescence microscopy (panel C). FITC-labeled anti-TREM2 antibody was used for the flow cytometry; typical flow cytometry data and the summary from 4 experiments are provided. Immunofluorescence microscopy results representative of at least 3 different experiments are shown. Rabbit anti-TREM2 primary antibody and Alexa 488-labeled secondary antibody were used to detect TREM2 expression in Meg-01 cells. **D & E.** Silencing CHOP reverses ER stress-mediated downregulation of TREM2 in Meg-01 cells. Typical blots (panel D) and their relative densities from 4 experiments (Supplemental Figure 3B) are shown. Panel E shows the typical immunofluorescence microscopy of TREM2 and CHOP expression in Meg-01 cells from at least 3 independent experiments. Meg-01 cells were infected with lentiviral CHOP-shRNA or control-shRNA, and then treated with tunicamycin 1 μg/mL for the indicated time. Rabbit anti-TREM2 primary antibody and Alexa-594 labeled secondary antibody were used to detect TREM2, and mouse anti-CHOP primary antibody and Alexa-488 labeled secondary antibody were used to detect CHOP. **F.** Intravenous injection of tunicamycin 1 mg/kg upregulates GRP78 and CHOP, and downregulates TREM2 without affecting C/EBPα in mouse megakaryocytes on day 3. Typical blots and their relative densities from 4 experiments are shown. **G & H.** Downregulation of TREM2 and upregulation of CHOP and GRP78 in a mouse model of MI. Mouse megakaryocytes and platelets were isolated from mice on day 3 and 7 after LAD ligation, respectively. Typical blots of megakaryocytes **(G)** and their relative densities from 6 experiments are shown. Typical blots of platelets **(H)** and their relative densities from 4 experiments are shown. **I.** Schematic diagram of TREM2 downregulation by CHOP-C/EBPα during ER stress: ER stress induces the upregulation of CHOP, which dimerizes with C/EBPα (a transcription factor of TREM2) to prevent its binding to TREM2 promoter. Decreased binding of C/EBPα to TREM2 promoter reduces TREM2 transcription and expression in megakaryocytes and further in platelets. Unpaired t-test was performed in panel F, G, and H.

Myocardial infarction (MI) is a serious event associated with CAD that usually occurs when the coronary arteries are severely blocked. During this process, excessive ER stress induces CHOP production and triggers cardiomyocyte apoptosis^14^. We next investigated the pathophysiological relevance of these observations using a mouse MI model. Consistent with our data from the tunicamycin-treated mice, MI increased CHOP and GRP78 levels in mouse megakaryocytes and platelets, indicating excessive ER stress, which subsequently caused a decrease in TREM2 level (Figure 3G and H). Collectively, these data support that ER stress downregulates TREM2 expression by the CHOP-C/EBPα axis (Figure 3I).

### TREM2 deficiency enhances mouse platelet aggregation, ATP release, P-selectin release, αIIbβ3 activation, and spreading

Considering the downregulation of TREM2 in the platelets from CAD patients, we investigated the function of TREM2 in platelets using TREM2-deficient mice. We found that TREM2 deficiency (Supplemental Figure 4A and B) slightly increased platelet count and volume (Supplemental Table VII) and had no effect on the number of α and dense granules in the platelets (Supplemental Figure 4C and D). TREM2 deficiency enhanced platelet aggregation in response to ADP, collagen, and CRP, and ATP release induced by collagen and CRP was also increased (Figure 4A, Supplemental Figure 5A - C). In contrast, platelet activation in response to thrombin or PAR4 receptor agonist AYPGKF was not affected by TREM2 deficiency (Supplemental Figure 6A and 6B). TREM2 deficiency also promoted P-selectin release and αIIbβ3 activation induced by ADP and CRP (Figure 4B).

**Figure 4.**
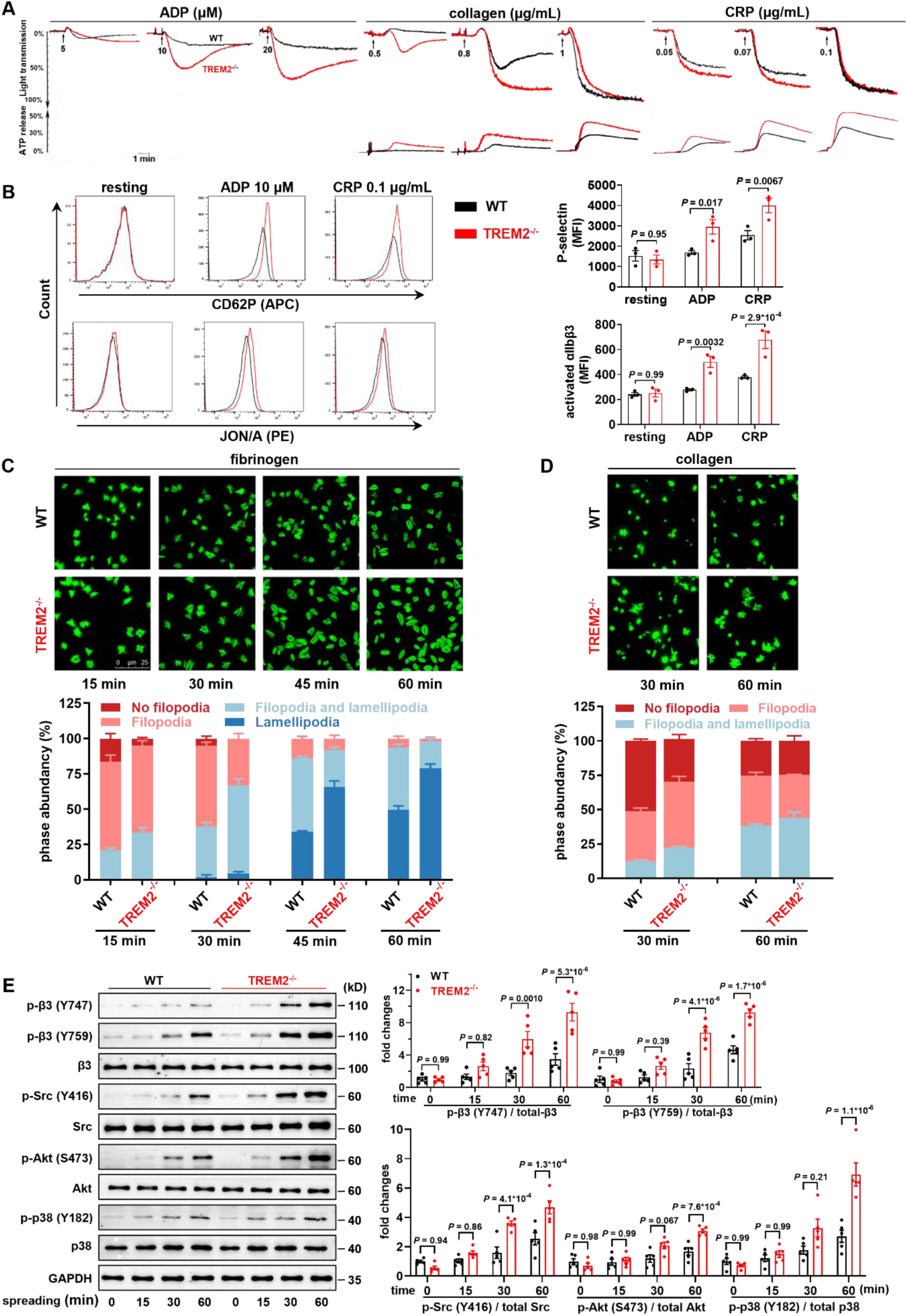
TREM2 deficiency enhances mouse platelet aggregation, ATP release, P-selectin release, αIIbβ3 activation, and spreading. **A.** TREM2 deficiency enhances mouse platelet aggregation in response to ADP, collagen, and CRP. ATP release induced by collagen and CRP is also enhanced. Washed platelets from wild-type (WT) and TREM2^-/-^ mice were stimulated with collagen (0.5, 0.8 or 1 μg/mL), CRP (0.05, 0.07 or 0.1 μg/mL) or ADP (5, 10 or 20 μM). The tracings are representative of 5 independent experiments using platelets from different mice. A summary of the data is provided in Supplemental Figure 5A and 5B. **B.** TREM2 deficiency enhances platelet P-selectin release and αIIbβ3 activation. ADP 10 μM, and CRP 0.1 μg/mL were used to activate platelets. APC-labeled anti-CD62P antibody and PE-labeled Jon/A antibody were used for flow cytometry. Typical histograms and the summary are provided. **C & D.** Typical photomicrographs of washed platelets from WT and TREM2^-/-^ mice spreading on immobilized fibrinogen (n = 4) and collagen (n = 3). Platelets were allowed to spread for the indicated time before being stained with phalloidin-FITC and photographed under fluorescence microscope. The phase percentage of these platelets was then calculated and shown as the mean ± SEM. **E**. TREM2 negatively regulates αIIbβ3 mediated outside-in signaling. WT or TREM2^-/-^ mouse platelets were plated on 20 μg/mL fibrinogen or 3% BSA as a negative control. Protein was extracted at 15, 30, and 60 minutes respectively and analyzed by Western blot. Typical blots and the summary of at least 5 independent experiments are provided. Two-way ANOVA followed by Sidak’s multiple comparisons test was used in **(B)**. Two-way ANOVA followed by Tukey’s multiple comparison test was performed in **(E)**.

Collagen-induced platelet aggregation is primarily mediated by glycoprotein VI (GPVI) which also contributes to platelet spreading by driving the formation of lamellipodial sheets and stress fibers^20^. Consistent with the enhanced platelet aggregation in response to collagen, TREM2^-/-^ platelets showed markedly enhanced spreading on fibrinogen and collagen compared with WT platelets, with faster formation of filopodia and lamellipodia (Figure 4C and D). TREM2 deficiency did not affect the thrombin-induced clot reactions (Supplemental Figure 6C), which is consistent with the unchanged platelet activation in response to thrombin in TREM2^-/-^ platelets (Supplemental Figure 6A and B).

### TREM2 deficiency enhances platelet outside-in signaling pathway

We further examined the effect of TREM2 deficiency on the outside-in signaling pathway during platelet spreading on immobilized fibrinogen. The results showed that the phosphorylation of β3 (Y747/Y759), c-Src (Y416), Akt (S473), and p38 (Y182) of TREM2^-/-^ platelets were significantly higher than WT, suggesting that TREM2 negatively regulates αIIbβ3-mediated outside-in signaling pathway (Figure 4E).

### TREM2 deficiency enhances platelet adhesion and thrombus formation

To further study the role of TREM2 in thrombosis, we used Bioflux microfluidic channels to mimic the hydrodynamic flow conditions and evaluate *ex vivo* thrombus formation in whole blood samples. TREM2^-/-^ platelets adhered faster and formed approximately 2.3 times larger thrombi than WT platelets at 5 min (Figure 5A).

**Figure 5.**
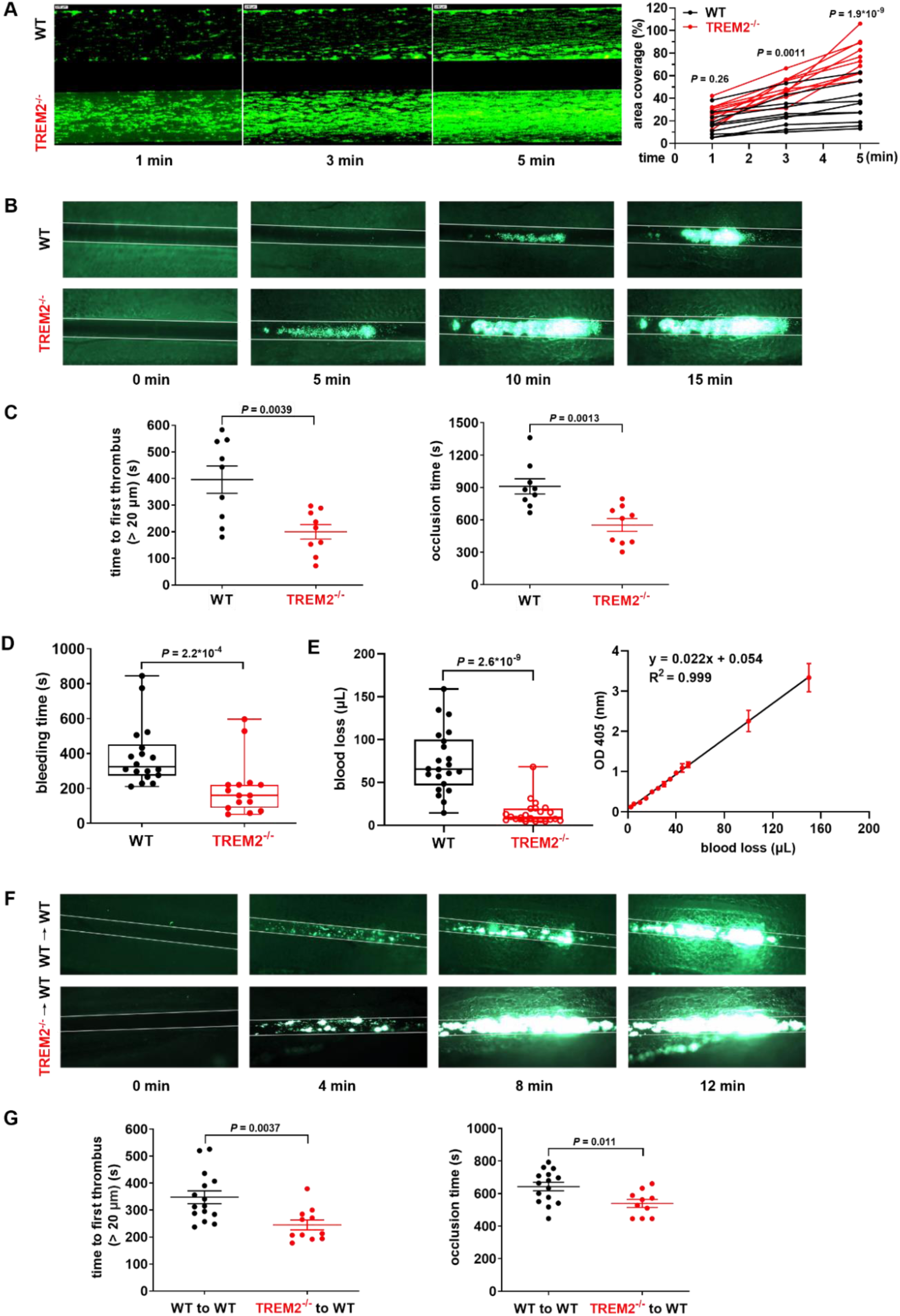
TREM2 deficiency promotes mouse platelet adhesion under flow conditions, accelerates thrombus formation, and reduces bleeding time and blood loss in mice. **A.** Quinacrine dihydrochloride labeled whole blood from WT and TREM2^-/-^ mice was perfused through a custom parallel plate flow chamber at a shear rate of 40 dynes/cm^2^ for 5 min. Platelet adhesion on collagen was monitored by fluorescence microscopy under a 10 ×lens. The percentage of chamber surface covered by platelets was analyzed by ImageJ (n = 10). Representative images of thrombus development over an immobilized collagen surface at different time points are shown. **B.** Typical images of accelerated thrombus formation in the FeCl_3_-injured mesenteric arterioles of TREM2^-/-^ mice. Calcein-AM was used to label platelets. **C.** Summary of panel B (n = 9). The occlusion time was defined by the stable occlusion for at least 2 min. **D & E.** TREM2 deficiency promotes mouse hemostasis. Reduced tail bleeding time in TREM2^-/-^ (n = 15) compared to WT (n = 18) mice **(D)**. Blood loss was measured after tail tip snipping using a well-established method. In the left panel, each dot represents one mouse (n = 22 in each group). The right panel is the standard curve used for blood loss assay **(E)**. **F.** Representative images showing increased thrombus formation in mesenteric arterioles of platelet-depleted WT mice repopulated with platelets from TREM2^-/-^ donors. **G.** Summary of panel F (n = 15 for WT, n = 11 for TREM2^-/-^). Two-way ANOVA followed by Sidak’s multiple comparisons test was performed in (A). Mann-Whitney test was performed in panel D and E. Data in panel C and G was analyzed by unpaired t-test.

Using the FeCl_3_-induced mesenteric arteriole thrombus formation assay, we found that TREM2 deficienct mice had increased thrombus formation in their mesenteric arterioles when compared to WT mice, as evidenced by the shorter time to first thrombus formation and arteriole occlusion (Figure 5B and C). TREM2^-/-^ mice also had a significantly shorter tail bleeding time (Figure 5D) and lower blood loss (Figure 5E) than WT mice suggesting a promoting impact of TREM2 deficiency on hemostasis.

To eliminate non-platelet hemostatic or thrombotic factor variations, we established a platelet depletion-reconstitution model by injecting 10^9^ platelets from TREM2^-/-^ or WT mice into thrombocytopenic WT mice^6,21,22^. Using FeCl_3_-induced thrombus formation assay, we found that infusion of TREM2^-/-^ platelets accelerates thrombus formation in mesenteric arterioles, as evidenced by the shorter time to first thrombus formation and arteriole occlusion (Figure 5F and G). This prothrombotic effect of TREM2 deficiency was independent of its role in platelet production, because the same number (10^9^ platelets) of WT and TREM2^-/-^ platelets were transfused into the recipient WT mice.

### TREM2 deficiency enhances inflammatory response in platelets and exacerbates experimental myocardial infarction

Platelets are important anucleate cells that link thrombosis and inflammation in cardiovascular disease^23,24^. TREM2 serves as an anti-inflammatory receptor in nucleated cells^25^. Thus, we investigated whether TREM2 influences the release of inflammatory cytokines from anucleate platelets. ELISA showed that TREM2 deficiency enhanced the release of proinflammatory cytokines, including IL-1β and TNFα, from activated platelets (Figure 6A), which are critical in the development of cardiovascular disease^26,27^. In contrast, TREM2 deficiency had no effect on the release of IL-6.

**Figure 6.**
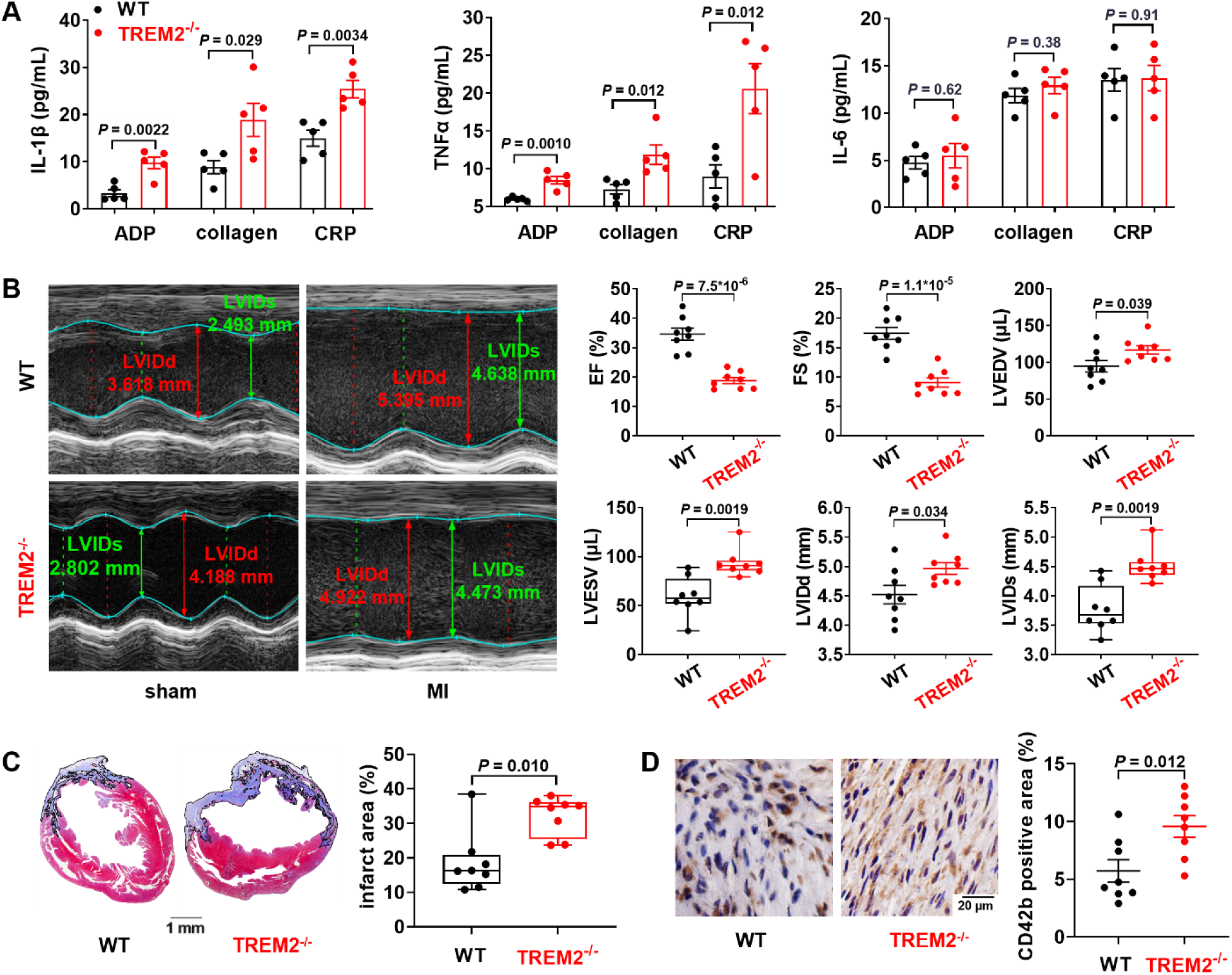
TREM2 deficiency promotes proinflammatory cytokine release from platelets and exacerbates experimental myocardial infarction (MI). **A.** TREM2 deficiency increases the release of IL-1β and TNFα, but not IL-6, from washed TREM2^-/-^ platelets stimulated by ADP (10 μM), collagen (0.8 μg/mL), or CRP (0.07 μg/mL), compared to washed wild type (WT) platelets, as determined by ELISA (n = 5). **B.** TREM2 deficiency exacerbates cardiac dysfunction. MI model was induced by left anterior descending coronary artery (LAD) ligation in WT and TREM2^-/-^ mice. M-mode echocardiography was taken on day 3 post-LAD ligation. Ejection fraction (EF), fraction shortening (FS), left ventricular end-diastolic/systolic volume (LVEDV/LVESV), and left ventricular internal diameter in diastole/systole (LVIDd/s) were calculated. Each dot represents a different mouse. Masson **(C)** and CD42b **(D)** staining of the hearts in panel B isolated on day 7 post-LAD. Unpaired t-test was performed in (**A**) and (**D**). Mann-Whitney test was performed in (**C**). LVESV and LVIDs in (**B**) were analyzed with Mann-Whitney test, while other data in (**B**) were analyzed with unpaired t-test.

Atherosclerotic plaque rupture induces excessive platelet activation and occlusive thrombosis in the coronary artery, which ultimately leads to MI. Ischemic stress following MI further activates platelets which form microthrombi in the myocardium and release proinflammatory cytokines, thereby exacerbating the myocardial infarction and reducing heart function^21,28–30^. After demonstrating that TREM2 deficiency enhanced platelet activation and inflammatory response, we further evaluated the contribution of these effects to experimental MI. Using a mouse MI model induced by left anterior descending coronary artery (LAD) ligation^21^, we found that TREM2 deficiency exacerbated cardiac dysfunction and increased the myocardial infarct area and microthrombus proportion in the peri-infarct area of these mice (Figure 6B - D).

### TREM2/DAP12/SHIP1 axis negatively regulate platelet activation via inhibiting Akt phosphorylation

TREM2 has a short intracellular domain that uses DNAX activating protein of 12 kDa (DAP12) as an adaptor to transduce intracellular signals^31^. As it is unclear whether platelets express DAP12, we showed that human platelets exhibit robust DAP12 expression at both the mRNA and protein levels (Supplemental Figure 7A and B), in line with DAP12 mRNA expression in human platelets^32^. Activation of macrophage TREM2 triggers the recruitment of DAP12 to TREM2, followed by the binding of Src homology 2 domain-containing inositol 5-phosphatase (SHIP1) to the TREM2/DAP12 complex in macrophages^33^. We explored whether the TREM2/DAP12/SHIP1 axis is also present in platelets. Immunoprecipitation assay showed that CRP treatment increased the recruitment of DAP12 to TREM2 and enhanced the binding of phosphorylated SHIP1 to the TREM2 complex in human platelets (Figure 7A), suggesting the presence of the TREM2/DAP12/SHIP1 axis in platelets. ADP, collagen, and CRP stimulation also increased phosphorylation of SHIP1 (Y1020, corresponding to human Y1022) and Akt (S473) in human platelets (Figure 7B).

**Figure 7.**
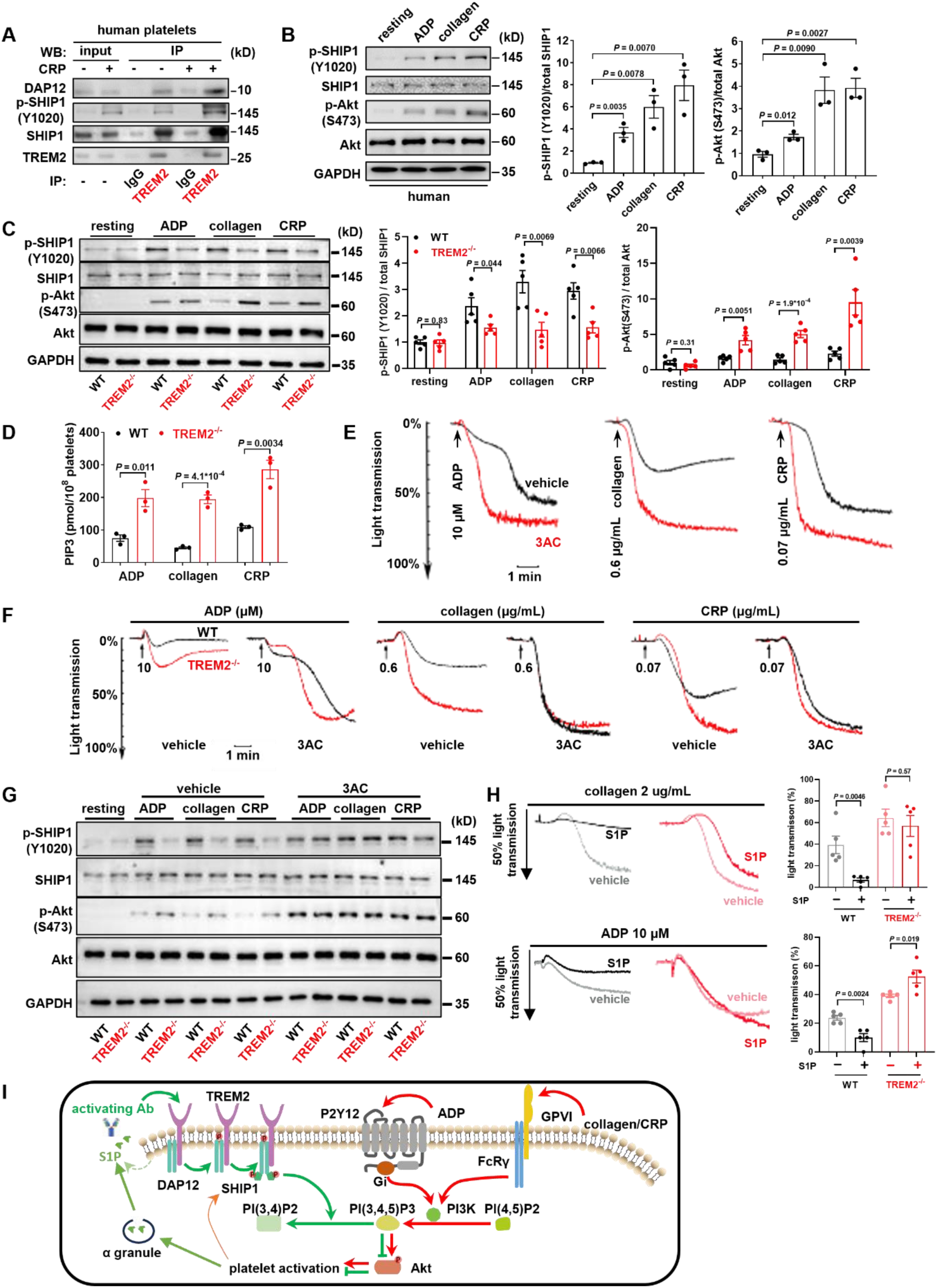
TREM2 negatively regulates platelet activation via DAP12/SHIP1/Akt downstream of GPVI and Gi. **A**. Increased DAP12, SHIP1, and p-SHIP1 co-immunoprecipitated with TREM2 in human platelets stimulated with 0.1 μg/mL CRP. Representative images of at least 3 independent experiments are shown. **B**. Increased p-SHIP1(Y1020) and p-Akt (S473) in human platelets stimulated with 10 μM ADP, 1 μg/mL collagen, and 0.1 μg/mL CRP. Representative blots and the summary of 3 independent experiments are shown. **C**. TREM2 deficiency reduces p-SHIP1 (Y1020), enhances the p-Akt (S473). Typical blots and the summary of at least 3 independent experiments are provided. **D**. TREM2 deficiency increases levels of phosphatidylinositol (3,4,5)-trisphosphate (PtdIns-(3,4,5)-P3, PIP3) in mouse platelets in response to 10 μM ADP, 0.8 μg/mL collagen, or 0.07 μg/mL CRP. The summary of at least 3 independent experiments is provided. **E**. SHIP1 inhibitor 3AC (10 μM) enhances the aggregation of washed human platelets. Typical tracings representative of three independent experiments and the summary of the data are provided. **F**. SHIP1 inhibitor 3AC (10 μM) potentiates platelet aggregation to a similar extent in WT and TREM2^-/-^ mice. Representative tracings of 3 independent experiments and the summary are shown. **G**. Similar phosphorylation levels of SHIP1 (Y1020) and Akt (S473) in activated WT and TREM2^-/-^ platelets in the presence of SHIP1 inhibitor 3AC. Samples from panel F were used for Western blot and representative blots and the summary from 3 independent experiments are shown**. H**. S1P inhibits platelet aggregation TREM2-dependently. S1P 5 μM inhibits platelet aggregation of wild type (WT) but not TREM2^-/-^ mice. Washed platelets were prepared from WT and TREM2^-/-^ mice, S1P 5 μM was used. Representative tracings of 5 independent experiments and the summary are shown. Data were present as mean ±SEM, unpaired t-test was used for statistical analysis. **I**. TREM2 regulates platelet activation via DAP12/SHIP1/Akt axis downstream of GPVI and Gi. S1P released from α granule during platelet activation is a TREM2 agonist that activates TREM2 to recruit and phosphorylate SHIP1 through DAP12. Phosphorylated SHIP1 converts PI(3,4,5)P3 to PI(3,4)P2, limiting Akt phosphorylation and platelet activation. Data in panel B and D were analyzed with unpaired t-test. Two-way ANOVA followed by Sidak’s multiple comparisons test was performed in panel C.

The TREM2/DAP12/SHIP1 axis is an important negative regulator of pro-inflammatory signals in nucleated cells^33,34^. Interestingly, recent studies have demonstrated that SHIP1 is a key negative regulator of platelet activation which acts through inhibiting Akt phosphorylation^35,36^. Upon platelet activation, SHIP1 is phosphorylated at Y1020^37^ and hydrolyzes phosphatidylinositol (3,4,5)-trisphosphate (PtdIns-(3,4,5)-P3, PIP3)^38^ to produce phosphatidylinositol (3,4)-bisphosphate (PtdIns-(3,4)-P2, PIP2), thereby reducing PIP3 levels. Reduced PIP3 attenuates Akt phosphorylation^39^ and provides important negative feedback preventing excessive platelet activation. Consistently, we found that TREM2 deficiency impaired the phosphorylation of SHIP1 (Figure 7C) and subsequently increased PIP3 levels (Figure 7D), thereby promoting Akt phosphorylation in agonist-treated mouse platelets (Figure 7C). Together, these results indicate that the TREM2/DAP12/SHIP1 axis functions in platelets to negatively regulate platelet activation via Akt inhibition.

We then investigated the role of SHIP1 in TREM2-mediated negative regulation of platelet activation using 3AC, a specific SHIP1 inhibitor^39–41^. Similar to TREM2 deficiency, treatment with 10 μM 3AC enhanced human platelet aggregation in response to ADP, collagen, and CRP (Figure 7E, Supplemental Figure 7C). Notably, 3AC treatment eliminated the differences of aggregation in response to ADP, collagen, and CRP between WT and TREM2^-/-^ platelets (Figure 7F, Supplemental Figure 7D). In line with the aggregation results, WT and TREM2^-/-^ platelets had similar phosphorylation patterns for SHIP1 and Akt in response to ADP, collagen, and CRP when they were treated with 3AC (Figure 7G, Supplemental Figure 7E). These results confirm that SHIP1 mediates the negative regulation of TREM2 on platelet activation (Figure 7I).

### S1P as an agonist of TREM2 inhibits platelet aggregation

Thus far, though we have shown that TREM2 is activated and inhibits platelet activation via DAP12/SHIP1 during platelet activation, the physiological ligand for TREM2 receptor is still not clear. Recently, Xue et al reported that S1P is an endogenous ligand of TREM2, and their functional study (microglia phagocytosis) indicates that S1P is a TREM2 agonist^42^. Platelets are also rich in S1P, which is released from α granules^43^, upon stimulation with collagen^44^, convulxin^43^, and ADP^43,45^. We proposed that upon platelet activation S1P is released from α granules, activates TREM2 receptor, and inhibits platelet activation. Consistently, we found that S1P inhibits platelet aggregation of wild type but not TREM2^-/-^ mice (Figure 7H), indicating that S1P inhibits platelet aggregation TREM2-dependently. Altogether, these data suggest that under physiological conditions, S1P released from platelet α granules is a physiological ligand which activates TREM2 receptor and restrain platelet activation.

### TREM2-activating antibody inhibits platelet aggregation TREM2 dependently

Given that TREM2 deficiency enhances platelet activation, we assumed that activation of the TREM2 receptor may inhibit platelet activation. A previous study demonstrated the feasibility of activating TREM2 using specific antibodies^46^. Therefore, we screened the commercially available antibodies against TREM2 and identified a TREM2-activating monoclonal antibody targeting the amino-terminus of mouse TREM2 (Supplemental Figure 8A). This TREM2-activating antibody inhibited the aggregation of WT, but not TREM2^-/-^ platelets induced by ADP, collagen, and CRP (Figure 8A and Supplemental Figure 8B).

**Figure 8.**
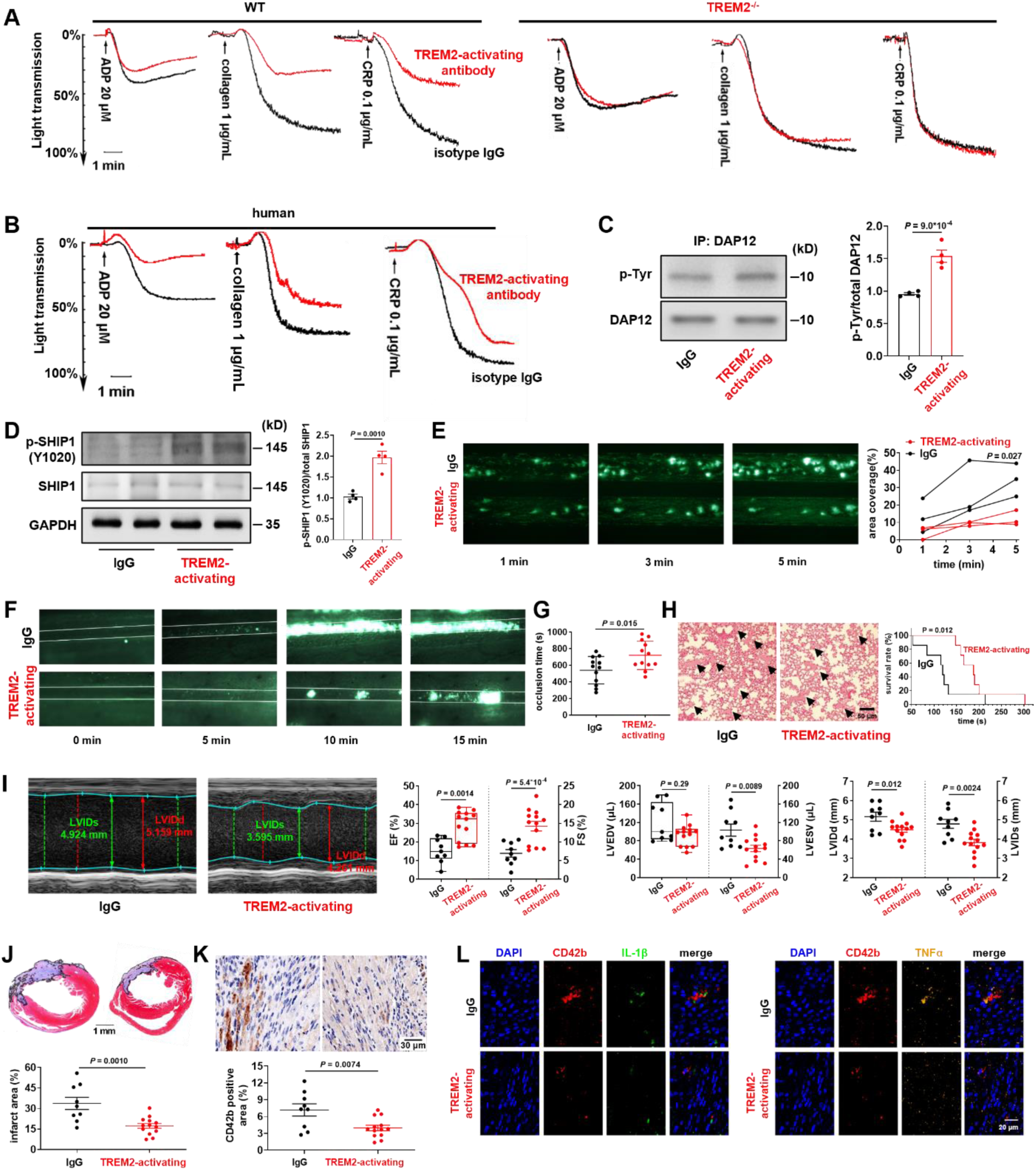
TREM2-activating antibody inhibits platelet aggregation and adhesion, and alleviates thrombus formation, pulmonary embolism, and experimental myocardial infarction in mice. **A.** The TREM2-activating antibody inhibits WT but not TREM2^-/-^ platelet activation in response to ADP, collagen, and CRP. **B.** The TREM2-activating antibody inhibits human platelet activation in response to ADP, collagen and CRP compared to IgG isotype. Washed platelets were pretreated with 12 μg/mL TREM2-activating antibody for 5 minutes before ADP (20 μM), collagen (1 μg/mL), and CRP (0.1 μg/mL) stimulation. IgG isotype was used as a control. A summary of the data is provided in Supplemental Figure 8B and C. **C & D.** The TREM2-activating antibody induces tyrosine phosphorylation of DAP12 and SHIP1. Human washed platelets were incubated with 12 μg/mL TREM2-activating antibody or lgG for 5 min at 37℃, then lysed, and immunoblotted to detect p-SHIP1. To detect tyrosine phosphorylation of DAP12, DAP12 was first immunoprecipitated followed by tyrosine phosphorylation immunoblot with phosphorpho-Tyrosine Mouse mAb (p-Tyr-100). Representative blots and the summary are shown (n = 4). **E.** Whole blood from C57BL/6 mice preincubated with 12 μg/mL TREM2-activating antibody or IgG isotype for 5 min was perfused through flow chamber coated with collagen at a shear rate of 40 dynes/cm^2^ for 5 min. Typical micrographs and a summary of the data are provided (n = 3). **F & G.** The TREM2-activating antibody inhibits FeCl_3_-induced thrombosis in mouse mesenteric arterioles. The TREM2-activating antibody or IgG (0.5 mg/kg) was intraperitoneally injected into mice 15 min before inducing thrombosis. Representative pictures and a summary of the data are provided (n = 12). **H.** The TREM2-activating antibody alleviates pulmonary thromboembolism and increases survival rate of mice with pulmonary thromboembolism. The TREM2-activating antibody (0.5 mg/kg) or IgG (0.5 mg/kg) was intraperitoneally injected into mice 15 min before pulmonary thromboembolism was induced by injection of a mixture of 430 μg/kg collagen and 20 μg/kg epinephrine. Survival rates were estimated using a Kaplan-Meier analysis and compared using the Breslow test (n = 9 for TREM2-activating antibody; n = 8 for IgG). **I.** TREM2-activating antibody injection improves cardiac function post-left anterior descending coronary artery (LAD) ligation. The TREM2-activating antibody or IgG used as a control were injected intraperitoneally at 0.5 mg/kg 15 min before LAD ligation to induce myocardial infarction. Echocardiography was carried out on day 3 post-LAD and typical echocardiographs and a summary of the data are provided (n = 13 for TREM2-activating antibody; n = 9 for IgG). Masson **(J)** and CD42b **(K)** staining of the heart tissues from panel I isolated on day 7 post-LAD ligation. Typical staining and the summary are provided. **L.** TREM2-activating antibody treatment reduces proinflammatory cytokines TNFα and IL-1β in the area corresponding to the infarct area in panel **J**. Successive slices of the mouse hearts isolated on day 7 post-LAD were prepared and subjected to immunofluorescent staining with antibodies for platelet CD42b (red), IL-1β (green), and TNF-α (orange); DAPI was used for cell DNA (blue). Typical micropgraphs shown are representative of 3 independent experiments using different mice. Unpaired t-test was used in (C), (D), (G), (J), and (K). Gehan-Breslow-Wilcoxon test was used in (H). Two-way ANOVA followed by Sidak’s multiple comparisons test was performed in (E). EF and LVEDV in (I) were analyzed with Mann-Whitney test, while other data in (I) were analyzed with unpaired t-test.

Consistently, the TREM2-activating antibody also inhibited human platelet aggregation stimulated by ADP, collagen, and CRP (Figure 8B and Supplemental Figure 8C). Moreover, the TREM2-activating antibody induced tyrosine phosphorylation of DAP12 and SHIP1 in human platelets (Figure 8C and D), further confirming its agonistic activity for TREM2.

### TREM2-activating antibody inhibits platelet adhesion and thrombus formation

We evaluated the *ex vivo* antithrombotic effect of the TREM2-activating antibody in whole blood using Bioflux microfluidic channels. The TREM2-activating antibody reduced human platelet adhesion by approximately 65.7% under flow conditions at 5 min (Figure 8E). We then explored the *in vivo* antithrombotic activity of the TREM2-activating antibody using an FeCl_3_-induced mesenteric arteriole thrombus model and a mouse pulmonary embolism model. Intravenous administration of the TREM2-activating antibody drastically slowed down thrombus formation in mouse mesenteric arterioles, with prolonged arteriole occlusion time (Figure 8F and G). The TREM2-activating antibody also alleviated pulmonary embolism and increased in median survival time (Figure 8H).

### TREM2-activating antibody inhibits experimental MI

Given our results that the TREM2-activating antibody suppressed platelet activation and thrombus formation, we next investigated its effects in the experimental MI model. Intravenous administration of the TREM2-activating antibody improved heart function, decreased infarct size (Figure 8I and J), and reduced the microthrombus formation measured by CD42b staining in mouse hearts following LAD ligation (Figure 8K). The TREM2-activating antibody reduced the inflammatory burden in hearts, as evidenced by the reduced presence of proinflammatory cytokines, IL-1β and TNFα which may also come from the nucleated cells, in the peri-infarct areas (Figure 8L). These data suggest TREM2-activating antibody might be an effective agent for treating MI-induced thrombosis and cardiac injury.

## Discussion

In this study, we demonstrated that the pattern recognition receptor TREM2 is an important node linking ER stress, innate immunity, and platelet hyperactivity: (1) platelets express functional TREM2; (2) TREM2 expression is decreased in platelets from patients with CAD, and this decrease is associated with platelet hyperactivity; (3) excessive ER stress downregulates platelet TREM2 via CHOP-C/EBPα; (4) TREM2 deficiency enhances platelet activation, *in vivo* thrombosis, and infract expansion post-MI; (5) TREM2/DAP12/SHIP1 axis negatively regulates platelet activation via Akt inhibition; (6) TREM2-activating antibody inhibits platelet activation, thrombosis, and protects against infract expansion post-MI in mice. Collectively, these results provide novel insights into the mechanisms underlying platelet hyperreactivity in patients with CAD: excessive ER stress in CAD impairs TREM2 expression in platelets, thereby induces platelet hyperreactivity, promotes thrombosis and infract expansion post-MI, and ultimately deteriorates CAD. Activating TREM2, for instance, using a TREM2-activating antibody, can serve as a novel antiplatelet strategy for patients with CAD.

The ER is one of the largest organelles, where approximately one-third of intracellular protein synthesis occurs. The oxidative environment within the ER promotes the formation of disulfide bonds that enable secretory and transmembrane protein folding. Disruptions in the folding capacity of ER proteins lead to a buildup of unfolded and misfolded proteins, and consequently result in the perturbation of ER homeostasis which is referred to as ER stress. We found that platelets from patients with CAD exhibit excessive ER stress, which induces platelet hyperactivity via CHOP-C/EBPα-mediated transcriptional repression of TREM2. Supporting our finding, previous study has shown that increased ER stress in platelets derived from diabetic animals contributes to enhanced thrombosis^47^. In addition, ER stress, which has been reported to increase CHOP^14,19,48^, is both a consequence and a cause of CAD. Notably, elevated CHOP levels in human arterial plaques maintained arterial plaque integrity^18,46^. When crossed with Apoe^-/-^ or Ldlr^-/-^ mice, CHOP-deficient mice have reduced growth, apoptosis, and necrosis in atherosclerotic plaques^49^, which support the notion that ER stress contribute to the aetiology of atherosclerosis. Our study further expands this notion and suggests that CHOP may also promote pathological arterial thrombus formation by downregulating TREM2, a negative regulator of platelet activation.

TREM2 is an innate immune receptor consisting of an extracellular immunoglobulin domain, a single transmembrane helix, and a short cytosolic tail^12,13,50^. Upon ligand stimulation, TREM2 associates with intracellular adaptor protein DAP12 and promotes its phosphorylation^12^. DAP12 contains an immune receptor tyrosine-based activation motif (ITAM), and through this ITAM, phosphorylated DAP12 recruits SH2 domain-containing protein tyrosine phosphatase SHIP1^34^. SHIP1 is a negative regulator of GPVI-mediated platelet activation^51^. The interaction of phosphorylated SHIP1 with PKCδ enhances its membrane localization, thereby increasing the catalytic activity of SHIP1 and leading to a decrease in PIP3 levels and Akt phosphorylation^37^. Akt phosphorylation regulates both integrin inside-out and outside-in signaling^52^. Platelet SHIP1 is phosphorylated and associated with PKCδ upon stimulation with GPVI agonist, while such association did not occur with PAR agonist. SHIP1 deficiency enhanced GPVI agonist-induced, but not PAR agonist-induced platelet aggregation^37^. Consistently, we found that TREM2 deficiency potentiates platelet activation induced by GPVI agonists collagen and CRP but not that induced by PAR agonists thrombin and AYPGKF. In addition, we found that TREM2 deficiency impairs the phosphorylation of SHIP1, and in turn relieves the SHIP1-mediated inhibition of PI3K/Akt by increasing PIP3 levels in platelets, while TREM2 activation has the opposite effect. Moreover, we demonstrated that TREM2 deficiency enhances inflammatory response in platelets, while TREM2 activation reduced the inflammatory burden. Recently, TREM2 expression was found in macrophages from both murine and human atherosclerotic lesions^53,54^, where TREM2 exerts an anti-inflammatory effect. These results, together with our findings on the antiplatelet and anti-inflammatory role of TREM2, indicate that TREM2 could be an attractive antithrombotic target in atherothrombotic diseases.

TREM2 is a new therapeutic target for tumor immune checkpoint therapy^55^ and Alzheimer’s disease^56,57^, and recent studies have identified TREM2 as a new target for atherosclerosis, with the TREM2-activating antibody 4D9 against atherosclerotic disease^54^. To our knowledge, the specific TREM2 ligand is still limited. Targeting the TREM2’s active domain with an activating antibody is now the most advanced therapeutic approach for TREM2. Currently, the activating monoclonal antibody against TREM2 (AL002) is tested in a phase II trial (NCT04592874) for early Alzheimer’s disease. And antibody PY314 is tested for solid malignancies (NCT04691375). In this study, we demonstrate that TREM2 activation by a TREM2-activating antibody inhibits platelet aggregation, secretion, and adhesion *in vitro*. In addition, intravenous administration of the TREM2-activating antibody suppresses experimental arterial thrombosis, alleviates pulmonary embolism, and improves heart function following LAD ligation *in vivo*, suggesting the potential of using TREM2-activating antibodies as antiplatelet and antithrombotic drugs. We also found that ursodeoxycholic acid (UDCA), a small molecular drug widely used for primary biliary cholangitis, also inhibits platelet activation and thrombosis as a TREM2 agonist (data not shown); TREM2 agonism may be a novel safe and efficacious approach for ASCVD treatment.

In conclusion, we demonstrated that excessive ER stress in CAD patients decreases platelet TREM2 expression via CHOP-C/EBPα mediated transcriptional suppression. Reduced TREM2 enhances platelet activation and accelerates thrombosis through the DAP12/SHIP1/Akt axis. We also provided evidence supporting that S1P is a physiological agonist of TREM2. Consequently, these signaling cascades aggravate infract expansion post-MI and heart dysfunction. Importantly, activation of TREM2 by TREM2-activating antibody may represent a novel strategy for decreasing the thrombotic burden in atherosclerotic diseases.

## Acknowledgments

We thank Drs. Xiaojing Gan and Liang Guo from the Fudan University School of Basic Medical Sciences for providing the pcDNA3.0-CHOP plasmid and pcDNA3.0-C/EBPα plasmid, respectively.

The Visual Abstract was modified from Servier Medical Art (http://smart.servier.com/), licensed under a Creative Common Attribution 3.0 Generic License.

## Funding

This work was supported by the National Natural Science of Foundation of China (Grant numbers: 81673429, 81872862, and 82173818), the Program for Professor of Special Appointments (Eastern Scholar) at Shanghai Institutions of Higher Learning, and the Startup Fund from Tianjin Medical University awarded to ZD.

## Authorship

XW, GP, LC, SZ and ZD designed research; XW, LC, GP, YL, YL, WZ, GZ, JZ, and HZ performed research; RX, YG, and ZQ contributed with patients and clinical data; SC, HH, JD, SZ, HH, and ZD analyzed data; XW, LC, GP, HH, JD, SZ, and ZD wrote and reviewed the paper.

## Ethic statement

All participants provided written informed consent to participate in the study. The study was approved by the Institutional Review Board of Fudan University. The research conforms with the principles of the Declaration of Helsinki.

## Conflict-of-interest disclosure

Authors declare no conflicts of interest.

## Notes

### Competing Interest Statement

The authors have declared no competing interest.

