## Supplemental Data for "ER stress-induced TREM2 downregulation exacerbates platelet activation and myocardial infarction in patients with coronary artery disease"

### Supplemental Material

#### Expanded Methods

##### Reagents

ADP, collagen, thrombin, and luciferin were purchased from Chrono-Log (Havertown, PA). Apyrase grade VII and human fibrinogen were purchased from Sigma-Aldrich (St. Louis, MO). PAR-4 activating hexapeptide (AYPGKF-amide) was synthesized by Shanghai Biotech BioScience & Technology (Shanghai, China) and Protein A/G Plus agarose was purchased from Santa Cruz Biotechnology (Santa Cruz, CA). The SHIP1 inhibitor, 3AC, was purchased from Minneapolis (Minneapolis, MN).

##### Human blood samples

Blood from healthy subjects and patients with CAD, including stable angina pectoris (SAP) and acute coronary syndrome (ACS), were obtained from the Fudan University Zhongshan Hospital. CAD was diagnosed using coronary angiography and informed consent was obtained. Basic clinical data for these participants are summarized in Supplemental Table III - V.

##### Platelet preparation

Human and murine platelets were isolated according to the previously described protocols[1-3]. Human blood samples were collected from the antecubital vein and anticoagulated with acid-citrate dextrose (85 mM sodium citrate, 71.38 mM citric acid, and 27.78 mM glucose, 6:1 vol/vol). For 20 mL of whole blood, platelet rich plasma (PRP) was obtained after 14 min of centrifugation at 300 × g. The PRP was centrifuged at 900 × g for 7 min and the palleted platelets were resuspended in Tyrode's buffer (138 mM NaCl, 2.7 mM KCl, 2 mM $MgCl_2$, 0.42 mM $NaH_2PO_4$, 5 mM glucose, 10 mM HEPES and 0.02 unit/mL apyrase, pH 7.4).

Mouse blood samples were collected from abdominal aorta and anticoagulated with 3.8% citrate (9:1). For 2 mL of whole blood, PRP was obtained after 2 min of centrifugation at 300 × g. The PRP was centrifuged at 450 × g for 2 min and the palleted platelets were resuspended in Tyrode's buffer.

##### Reverse transcription polymerase chain reaction (RT-PCR) and qPCR

Total RNA was extracted using TRIzol Reagent (Invitrogen, NY) and reverse transcribed to cDNA using an RT-PCR kit (TaKaRa, Dalian, China). qPCR was performed using specific primers (Supplemental Table I) and TB Green Premix Ex Taq II (TaKaRa, Dalian, China) according to the manufacturer's protocol on an Applied Biosystems 7500 qPCR System (Thermo

Fisher, Shanghai, China).

**Western blot**

Freshly prepared platelets ($3 \times 10^8$ /mL) were stimulated with the indicated agonist for 3 min. Reactions were then stopped by adding 5× loading buffer (Beyotime Biotechnology, Shanghai, China), boiled for 5 min, and subjected to Western blot analysis. The antibodies used are listed in Supplemental Table II.

**Immunofluorescence staining**

Platelets resuspended in Tyrode's buffer ($5 \times 10^7$ /mL) were fixed and permeabilized using Cytofix/Cytoperm™ Fixation/Permeabilization solution (Becton Dickinson and Company, NY) at room temperature for 15 min, then incubated with the appropriate primary antibodies overnight at 4°C, followed by incubation with Alexa Fluor-594-labeled or Alexa Fluor-488-labeled secondary antibodies for 2 h at room temperature. The slides were washed and then placed in the dark at 4°C until image acquisition.

For immunofluorescence microscopy of the Meg-01 cells, the procedure was similar to that described above. Briefly, Meg-01 cells were quickly spun, resuspended in a minimal volume of 2% PFA/PBS, and spread evenly on a coverslip. After incubating with the appropriate primary antibodies overnight at 4°C, these coverslips were then incubated with diluted Alexa-conjugated secondary antibodies combined with 100 ng/mL DAPI for 1 h in the dark at room temperature. Fluorescent signals were detected using a confocal laser scanning microscope (SP8, Leica, Germany).

**Flow cytometry**

Blood samples from patients with ACS were collected before they received any antiplatelet therapy. Platelets were separated and incubated with the indicated agonists at 37°C for 15 min before being incubated with the appropriate fluorophore-conjugated antibodies (P-selectin/TREM2, shown in Supplemental Table II) for 60 min at room temperature in the dark. Tyrode's buffer was added to each sample to terminate the reaction and the samples were immediately analyzed using a FACSCalibur flow cytometer (BD FACSCelesta, NJ).

Meg-01 cells stimulated with tunicamycin were centrifuged at 1000 rpm for 3 min and then washed with PBS before being fixed and permeabilized in Cytofix/Cytoperm™ Fixation/Permeabilization solution (Becton Dickinson and Company, NY). These cells were then resuspended in PBS and incubated with FITC-conjugated TREM2 antibody (Supplemental Table II) for 60 min at room temperature in the dark. The cells were washed with PBS and analyzed using a FACSCalibur flow cytometer (BD FACSCelesta, NJ).

**Cloning of different types of TREM2 promoters into a pGL3 reporter vector**

Human TREM2 promoters with different length and mutation were cloned into a pGL3 reporter vector for luciferase assay[4,5] and verified by sequencing. Briefly, a serial of truncated constructs of human TREM2 promoter 3.0 kb upstream from the transcriptional start site was amplified using genomic DNA extracted from normal HeLa cells by PCR using different primers (Supplemental Table I). Overlap-extension PCR was used to generate the deletion mutant of TREM2 promoter $^{-298}$TTGCA$^{-294}$. The deletion was confirmed via sequencing.

**Transient transfection and luciferase assay of TREM2 promoters**

Transient transfection and luciferase assay were performed as previously described. Firefly luciferase constructs containing the various TREM2 promoter fragments were transiently transfected into HEK293T cells using Lipo3000 (Invitrogen, NY) and assayed for promoter activity. Briefly, $6 \times 10^4$ cells were plated in 24-well plates 24 h before the experiment. The mixture of 2.4 μL Lipo3000 and 25 μL of Opti-MEM media were prepared, and then 0.4 μg of pGL3 plasmid containing the TREM2 promoter (wild-type, mutant, or truncated constructs) and the CHOP or C/EBPα plasmid were added and incubated for 20 min to form DNA-lipid complexes in the presence of 16 ng pRL-TK plasmid. The cells were then transfected with the mixture for 6 h, and then the culture medium was changed. The cells were cultured for an additional 48 h and then washed twice with cold PBS. Cell lysates were prepared by adding 100 μL of lysis buffer, and the protein supernatant was collected. Firefly and Renilla luciferase activities were assessed using the Dual-Glo luciferase assay system (Promega, WI) in accordance with the manufacturer’s instructions. Luminescence readings were acquired using a TD 20/20 luminometer (Turner Design Inc., Sunnyvale, CA) and sample values were compared to the reference value of pGL3-Basic empty vector/empty pRL-TK. At least three independent transfections were performed.

**Lentiviral coating and infection**

Lentivirus was constructed as previously described[5]. The shRNA against CHOP amplified by the relevant primer described in Supplemental Table II was subcloned into the PHY-310 vector between the *BamH1* (2440) and *EcoRI* (2457) restriction sites. The recombinant plasmid was then co-transfected with psPAX2 and pMD2.G (4:3:1) into HEK293T cells using lipofectamine 3000 (Invitrogen, NY) according to the manufacturer's instructions. After 6-hours culture, the medium was changed, and the cells were cultured for an additional 72 h. The culture media was centrifuged at 3000 rpm for 10 min and the virus-containing supernatant was filtered through a 0.45 μm filter. Lentivirus supernatant and polybrene (final concentration of 8 μg/mL) were mixed and then added to $6 \times 10^4$ cells/mL Meg-01 cultured in a 60 mm dish. After 6 h, the culture medium was changed, and the cells were cultured for a further 48 h. The infection efficiency was determined by cellular fluorescence.

**Electrophoretic mobility shift assay (EMSA)**

The EMSA was constructed as previously described[6]. The 5'-biotin-labeled (5'-CGGGCA**TTGCA**GCTGGTGGA-3'), unlabeled and mutant oligonucleotide probes (5'-CGGGCA**GCGCA**GCTGGTGGA-3') corresponding to the C/EBPα binding site in the TREM2 promoter region (-294 to -298) were double-stranded and purchased from Bioscience Technology (Shanghai, China). Sample response systems (total volume, 10 μL) were produced according to the manufacturer's instructions and the reaction solution included 10 mM Tris-HCl (pH 8.0), 10 mM $MgCl_2$, 1 mM EDTA, 1 mM dithiothreitol, 10% glycerol, and 60 mM KCl. Purified C/EBPα protein (5 μg) was used as a specific transcription factor and was inhibited by 5 μg of purified CHOP protein (Supplemental Figure II). In the probe competitive reaction, a 100-fold unlabeled probe was used. For supershift analysis, 2 μg of antibody (anti-C/EBPα or isotype lgG, Supplemental Figure II) was added to the incubation mixture at room temperature 1 h prior to the addition of the labeled probe.

The samples were incubated at room temperature for 20 min and then separated from free DNA probes using a 6.5% non-denaturing polyacrylamide gel in 0.5× TBE buffer at 10 V/cm and electrophoretically transferred onto a nylon membrane (Beyotime Biotechnology, Shanghai, China) for 30 min at 380 mA. The transferred DNA was cross-linked on a membrane illuminated with UV light for 20 min at 10 cm, and the membrane was blocked in 1× blocking buffer before being incubated with a streptavidin-HRP conjugate. After proper washing (5 times for 4 min), the signals were visualized using a chemiluminescence imaging system.

**Chromatin immunoprecipitation (ChIP) assay**

ChIP assay was performed using a ChIP Assay Kit (9005S, Cell Signaling Technology, MA) according to the manufacturer's protocol with minor modifications[2]. Meg-01 cells were grown to 90% - 95% confluence in a 10 cm dish and treated with 1% formaldehyde for 10 min at 37°C to cross-link the proteins and DNA, and then stopped with 0.125 M glycine. After washing three times with ice-cold PBS, the cells were scraped into a conical tube before being pelleted, lysed, sonicated, and immunoprecipitated using a C/EBPα antibody (Santa Cruz Biotechnology, CA) or isotype IgG (negative control). Immune complexes (except for 10 μL of supernatant saved as input) were collected using 50 μL of protein A agarose beads for 60 min at 4°C with rotation and washed once with low-salt wash buffer (0.1% SDS, 1% Triton X-100, 2 mmol/L EDTA, 20 mmol/L Tris-HCl (pH 8.0), 150 mmol/L NaCl), high-salt wash buffer (0.1% SDS, 1% Triton X-100, 2 mmol/L EDTA, 20 mmol/L Tris-HCl (pH 8.0), and 1.5 mol/L NaCl), and LiCl wash buffer (250 mmol/L LiCl, 1% NP-40, 1% sodium deoxycholate, 1 mmol/L EDTA, and 10 mmol/L Tris-HCl (pH 8.0)), and twice with 10 mmol/L Tris-HCl (pH 8.0) and 1 mmol/L EDTA. Immune complexes were eluted using freshly prepared elution buffer (1% SDS in 0.1 mol/L $NaHCO_3$) and cross-linking was reversed by heating at 65°C overnight in the presence of 0.2 M NaCl. TREM2 enrichment was evaluated by qPCR.

**Myocardial infarction model**

The mouse myocardial infarction model was established as previously described[1,7,8]. Mice were placed on a heating pad and anesthetized in an airtight chamber using 5% isoflurane. After endotracheal intubation, a left thoracotomy was performed in the second and third intercostal spaces. Using an 8-0 polypropylene suture (Ningbo Chenghe Micro Instrument Factory, Ningbo, China), the exposed heart was ligated at the proximal portion of left anterior descending artery. The chest and skin were then closed in layers using 6-0 silk sutures and the mice were extubated and warmed for several minutes to allow for recovery[9]. M-mode echocardiographic images were taken on day 3. Mouse hearts were collected on day 7 and sham operations were performed in the same way, except that ligation of the left anterior descending coronary artery was not performed to act as the control.

**Mouse bone marrow isolation and CD61$^+$ megakaryocyte preparation**

Mouse bone marrow aspirate samples were collected in tubes containing EDTA. CD61$^+$ megakaryocytes were isolated using CD61 MicroBeads (Miltenyi Biotech, Germany) according to the manufacturer's instructions as previously reported[2].

**Platelet aggregation and secretion**

Platelet aggregation and ATP release were assayed as described previously[1,10]. The washed platelets were adjusted to the concentration of $3 \times 10^8$ /mL and preincubated with the TREM2-activating antibody, IgG isotype or 3AC for 5 min. Then, in the present of luciferin, platelet aggregation and ATP release were triggered with agonists under stirring conditions (900 rpm) at 37°C and recorded simultaneously.

**Platelet spreading**

Platelet spreading was evaluated as described previously[1-3]. The washed platelets ($2 \times 10^7$ /mL) were allowed to spread on Lab-Tek chamber slides (Nalge Nunc International, Rochester, NY) precoated with 20 μg/mL of fibrinogen for indicated time at 37°C. After washing with PBS, attached platelets were fixed and permeabilized using Cytofix/Cytoperm™ Fixation/Permeabilization solution (Becton Dickinson and Company, NY), and then stained with FITC-labeled phalloidin. Fluorescent signals were detected using a confocal laser scanning microscope (SP8, Leica, Germany). Depending on activation status of platelets spread on immobilized fibrinogen, platelets 1) with no filopodia, 2) with filopodia, 3) with filopodia and lamellipodia, and 4) with lamellipodia will be observed as time goes on. Platelets with different shape were counted in 3 eye fields and calculated. Platelet phase abundance in each independent experiment was calculated from three eye fields.

**Clot retraction**

Clot retraction was evaluated as described previously[1-3]. Washed platelets ($3 \times 10^8$ /mL) mixed with 2 mg/mL fibrinogen were dispensed in 0.3 mL aliquots into cuvettes. Clot retraction was triggered by the addition of 1 U/mL thrombin and allowed to proceed at 37°C. Photographs were taken at the indicated time points. Sizes of clots were quantified using ImageJ software.

**Flow chamber assay of platelet adhesion**

The flow chamber assay of platelet adhesion was performed as described previously with minor modifications[11]. Briefly, platelet adhesion in flow chamber was evaluated using a microfluidic whole-blood perfusion system under arterial shear conditions in a Bioflux-200 system (Fluxion, South San Francisco, CA). Bioflux plates were coated with collagen (100 μg/mL) overnight, then blocked using 5% BSA for 1 h before the plates were examined under an inverted microscope (Nikon Ti-S, Tokyo, Japan). Mepacrine-labeled blood was then perfused through the inlet well at a shear rate of 40 dynes/cm$^2$ for 5 min and adhered platelets were viewed using an S Plan Fluor lens (× 20/0.4 numerical aperture objective). Images were acquired using a Nikon DS-Qi1-U3 CCD camera and the platelet-covered area was measured using ImageJ software.

**Bleeding time assay**

The mouse tail bleeding time was assayed as previously described[3]. Briefly, tails of the wild-type (WT) and TREM2$^{-/-}$ mice were transected 5 mm with surgical scissors. The tail was immersed in a 15 mL tube containing 14 mL saline prewarmed to 37°C and the time taken for the bleeding to stop (no rebleeding within 30 s) was recorded.

**Tail blood loss assay**

Tails of the WT and TREM2$^{-/-}$ mice were transected 5 mm with surgical blades. The tail was immediately immersed in 0.9% saline at 37°C. No bleeding within 30 seconds counts as bleeding stops. Normal saline mixed with blood was frozen and thawed at -80 degrees, red blood cell lysate was used to lyse red blood cells to release hemoglobin, and the absorbance of 405 nm was detected by a microplate reader. The volume of blood loss was assessed by measuring the amount of hemoglobin. At the same time, a standard curve was prepared, the abdominal aortic blood of C57BL/6J mice was taken, the whole blood was diluted with normal saline according to different dilution factors, a series of standards were prepared, the absorbance was detected at a wavelength of 405 nm on a microplate reader, and the standard curve was drawn after conversion.

**Intravital microscopy of $FeCl_3$-injured thrombus formation in murine mesenteric arterioles**

Intravital microscopy of $FeCl_3$-induced thrombus formation in mouse mesenteric arterioles was performed as previously described[1,3]. Briefly, calcein-labeled platelets were injected into WT or TREM2$^{-/-}$ mice via the tail vein and thrombosis was induced using 10% $FeCl_3$ for 5 min and recorded using intravital microscopy.

**Pulmonary thromboembolism mouse model induced by collagen and epinephrine**

C57BL/6J mice were intraperitoneally injected with 0.5 mg/kg TREM2-activating antibody or lgG isotype. After 15 min, pulmonary embolism was induced by tail vein injection of recombinant collagen at 430 μg/kg and epinephrine at 20 μg/kg, as described previously[12]. Death time was recorded over a 15 min period.

**Platelet depletion/reconstitution model**

As described before[1,3], platelet depletion/repletion was performed according to the method originally reported by Robins et al[13] with minor modification，a well-established method in our lab[1,3]. Briefly, receiver mice were injected with rabbit anti–mouse thrombocyte serum (20 µL) intraperitoneally (Cat. #: J1943, Accurate Chemical and Scientific Corporation). After 4 hours, blood was collected by tail tip cutting for platelet count using Auto Hematology Analyzer (Mindray BC-2800Vet, Shenzhen, China), platelet depletion was considered successful if the total level of circulating platelets decreased to less 100 × $10^9$/L. Then$10^9$ platelets were injected via the tail veins to reconstitute the platelets, before being subjected to an in vivo

FeCl3-induced thrombus formation assay and the MI model.

**Enzyme-linked immunosorbent assay of inflammatory cytokines in mouse platelets**

Washed platelets from mice were treated with ADP, collagen, or CRP for 4 h at 37°C. After centrifugation at 450 × g for 2 min, the supernatant was then collected, and the levels of IL-1β, IL-6, and TNFα were determined using commercial ELISA kits (Invitrogen, NY) according to the manufacturer's instructions.

**Immunoprecipitation**

Washed platelets were added to equal volumes of ice-cold 2× lysis buffer (300 mM NaCl, 10% 1 mM Tris-HCl (pH 7.4), 2% NP-40, 0.5 M EDTA, 1 mM NaF) containing a protease and phosphatase inhibitor cocktail. Platelet lysates were incubated with an antibody against TREM2 or isotype control overnight at 4°C before prewashed protein A/G plus-agarose was added, and the immune complexes were rotated for 3 h. Samples were then centrifuged at 12000 × g for 30 s to pellet the agarose beads, before being washed with lysis buffer and then boiled in reducing SDS-PAGE sample loading buffer for 3 min for Western blot analysis.

**Phosphatidylinositol (3,4,5)-trisphosphate (PtdIns-(3,4,5)-P3, PIP3) assay**

Washed mouse platelets in 1 mL Tyrode's buffer ($3 \times 10^8$ /mL) were incubated with the specified agonist at 37°C for 5 min and then centrifuged at 450 × g for 3 min, before the platelet pellets were resuspended in 5 mL cold 0.5 M TCA and incubated on ice for 5 min. PIP3 was then extracted and assayed using a PIP3 MASS ELISA kit (Echelon, IL) according to the manufacturer's protocol.

**Statistical analysis**

Statistical analyses were performed using GraphPad Prism 9. For data conforming to Gaussian distribution, unless otherwise stated, differences between the two groups were analyzed by unpaired Student t-test. One-way ANOVA followed by Tukey's test or two-way ANOVA followed by Sidak's test was used for multiple comparisons. For data that did not conform to Gaussian distribution, the non-parametric Mann-Whitney test was used for the comparisons of two groups and Kruskal-Wallis test followed by Dunn's test for multiple comparisons. Pearson's correlation was used to analyze the correlation between the two continuous variables. Gehan-Breslow-Wilcoxon test was used for the comparison of survival curves. Differences were considered significant at $P < 0.05$.

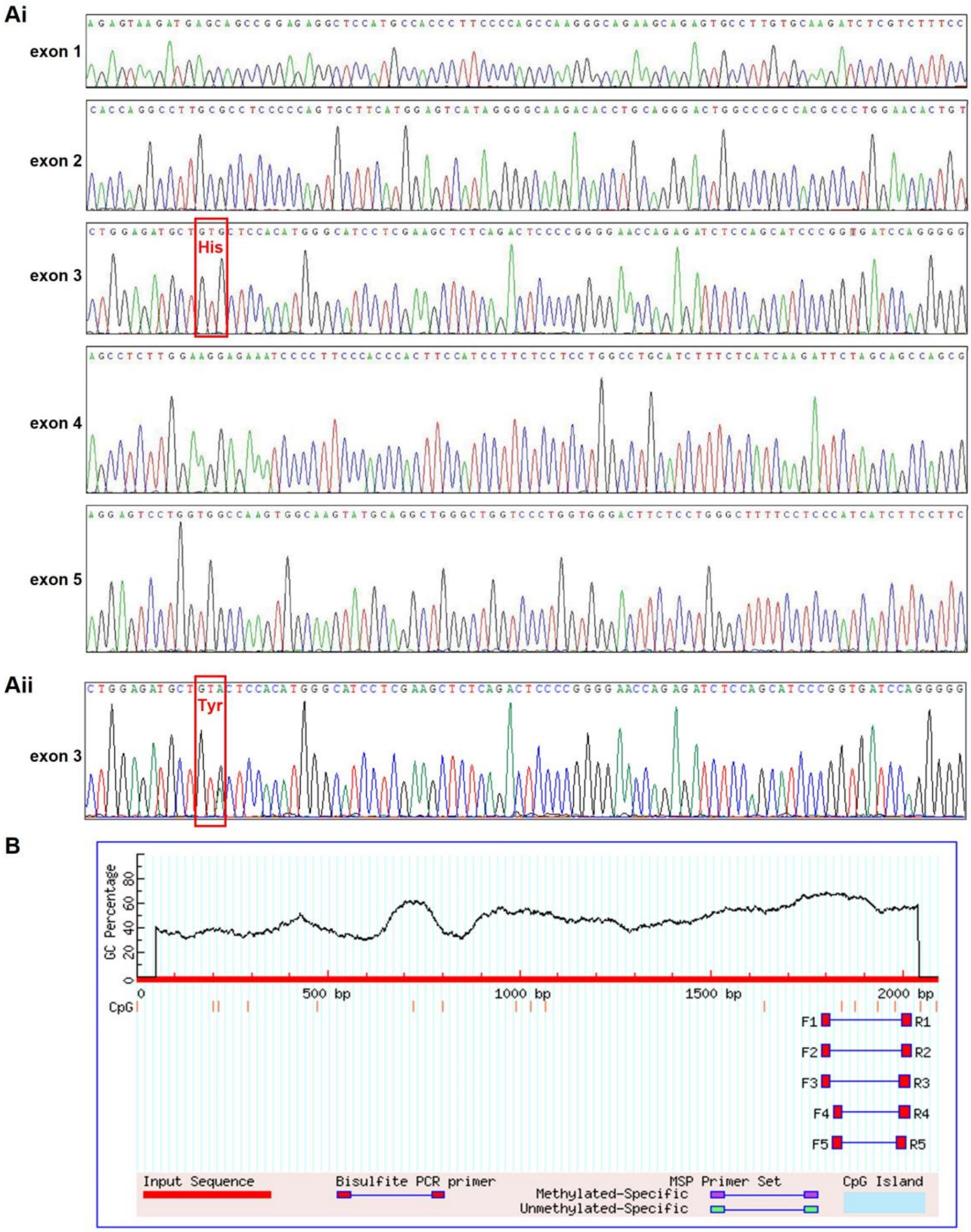


**Supplemental Figure 1. Genetic alterations and epigenetic inactivation are not responsible for TREM2 downregulation. A.** The sequence of TREM2 exons detected by Sanger sequencing in 50 ACS patients. **i)** Part of the reported reference sequence of all 5 exons of human TREM2 (GenBank accession number: NM_018965.3). **ii)** c.469G > A (p.His157Tyr) mutation (red frame) in exon 3 was identified in one of the 50 ACS patients. **B.** CpG island was not found in TREM2 promoter (-2000 - 0) sequence using MethPrimer (http: //www.urogene.org/methprimer). Characteristics of the study population were provided in Supplement Table VI.

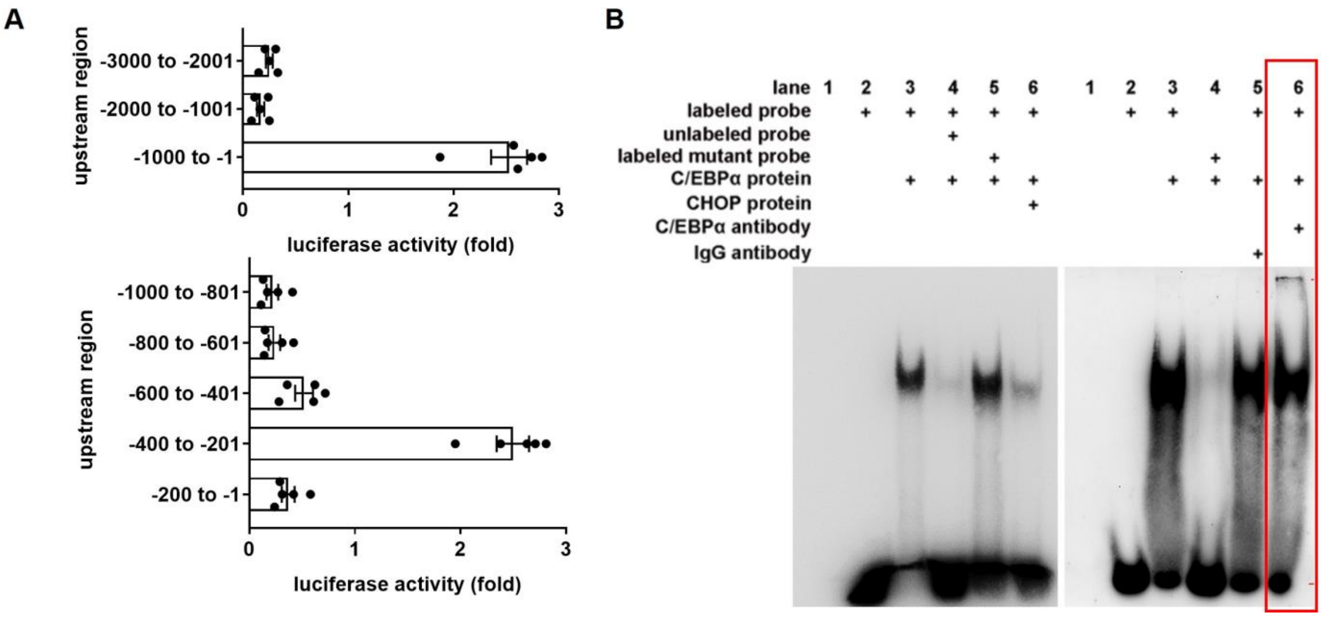


**Supplemental Figure 2. CHOP inhibits C/EBPα-mediated TREM2 transcription. A.** Transcriptional activities of truncated TREM2 promoters detected by luciferase reporter assay (n = 5). **B.** EMSA assay showing C/EBPα binds TREM2 promoter in Meg-01 cells, which is impaired by CHOP. Nuclear protein extracted from Meg-01 cells was incubated with biotin-labeled probe (5’-CGGGCA**TTGCA**GCTGGTGGA-3') containing C/EBPα-response element ($^{-298}$TTGCA$^{-294}$) in TREM2 promoter or mutated probe (5’-CGGGCA**GCGCA**GCTGGTGGA-3'). Left: the representative picture of electrophoretic mobility shift assay (EMSA). Right: EMSA antibody supershift assay with anti-C/EBPα antibody or IgG isotype. A DNA-protein complex band was detected when nuclear protein was incubated with a probe containing $^{-298}$TTGCA$^{-294}$ (left panel, lane 3) and this complex band was abolished in the presence of competing unlabeled oligonucleotides (left panel, lane 4) or CHOP protein (left panel, lane 6). There were no obvious DNA-protein complexes when a probe containing a mutated C/EBPα-response element was used (right panel, lane 4). The DNA-protein complexes were super-shifted in the presence of an anti-C/EBPα antibody (right panel, lane 6). Typical results representative of 3 independent experiments are shown.

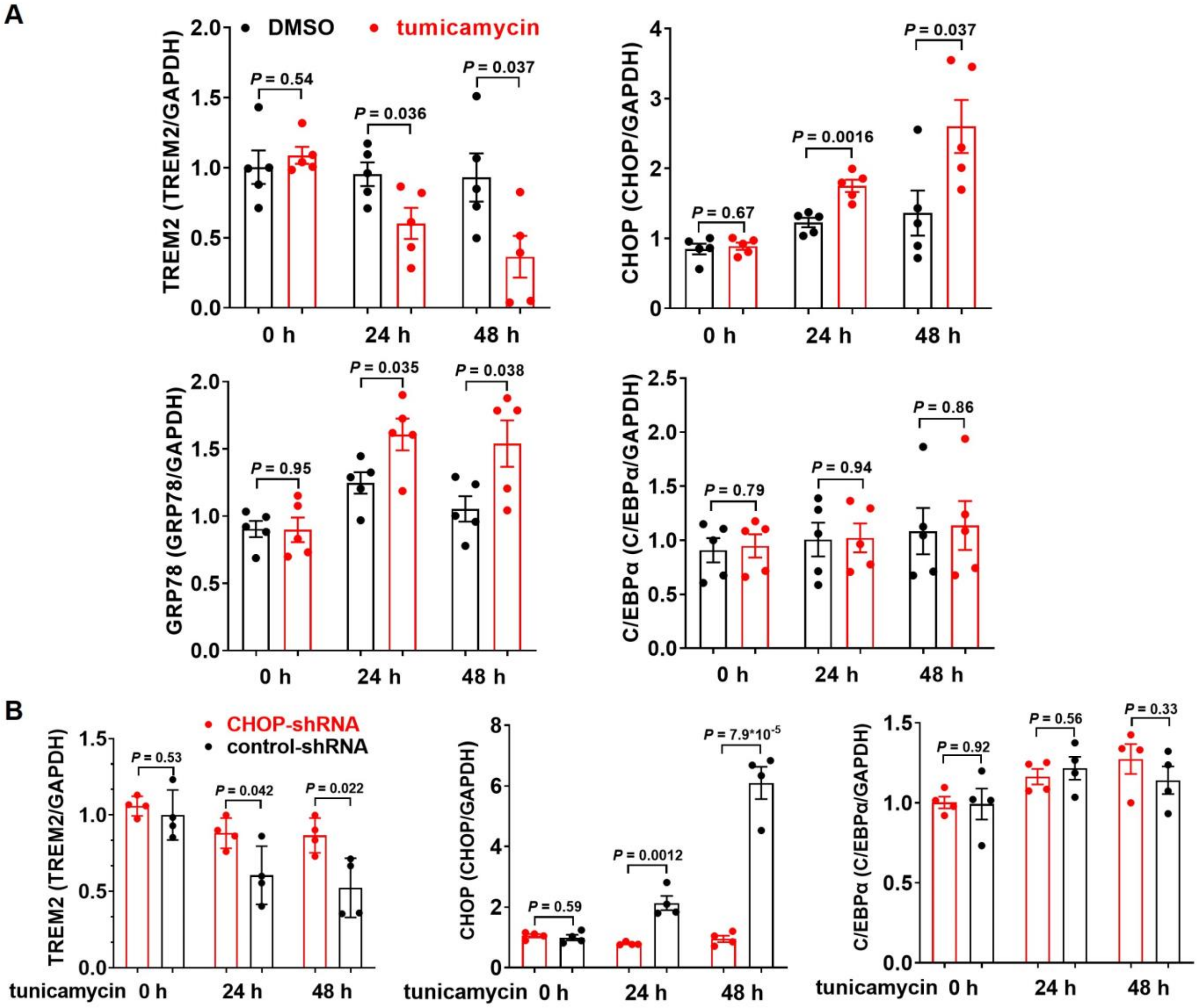


**Supplemental Figure 3. ER stress downregulates TREM2 expression in Meg-01 and platelets. A.** Tunicamycin upregulates GRP78 and CHOP, and downregulates TREM2 expression without affecting C/EBPα levels in Meg-01 cells. Relative densities from 5 experiments, corresponding to Figure 3A, are presented. **B.** Silencing CHOP reverses ER stress-mediated downregulation of TREM2 in Meg-01 cells. Relative densities from 4 experiments, corresponding to Figure 3D, are presented. Unpaired t-test was performed in panel A and B.

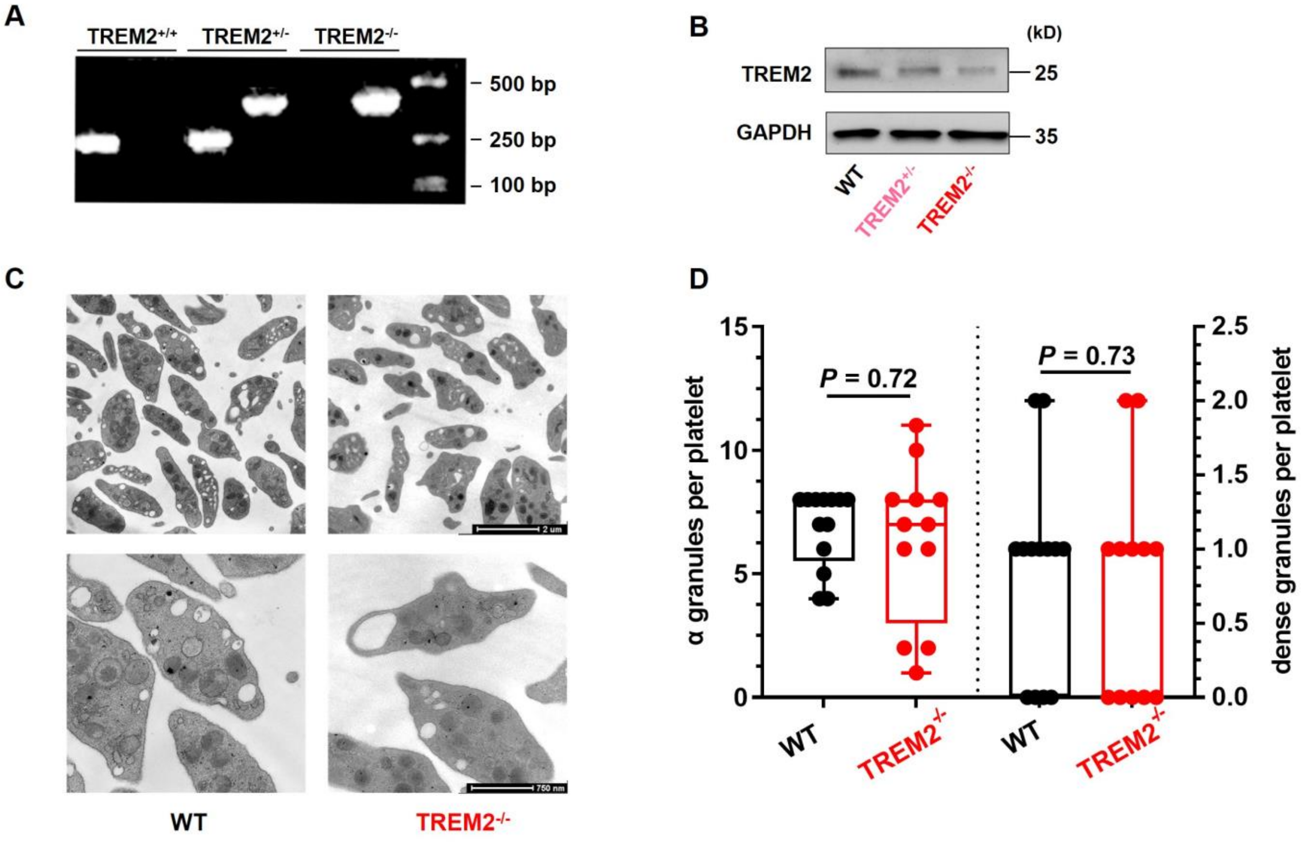


**Supplemental Figure 4. TREM2 deficiency does not affect α granules and dense granules in mouse platelets. A.** Genotyping of TREM2$^{-/-}$ mice. Wild-type (WT): 250 bp; TREM2$^{-/-}$: 396 bp. **B.** TREM2 expression was reduced in platelets of TREM2$^{-/-}$ mice. **C.** Transmission electron microscopic images of WT and TREM2$^{-/-}$ platelet ultrastructure. **D.** The counts of α granules and dense granules in WT (n = 13) and TREM2$^{-/-}$ (n = 12) platelets. Mann-Whitney test was performed in panel D.

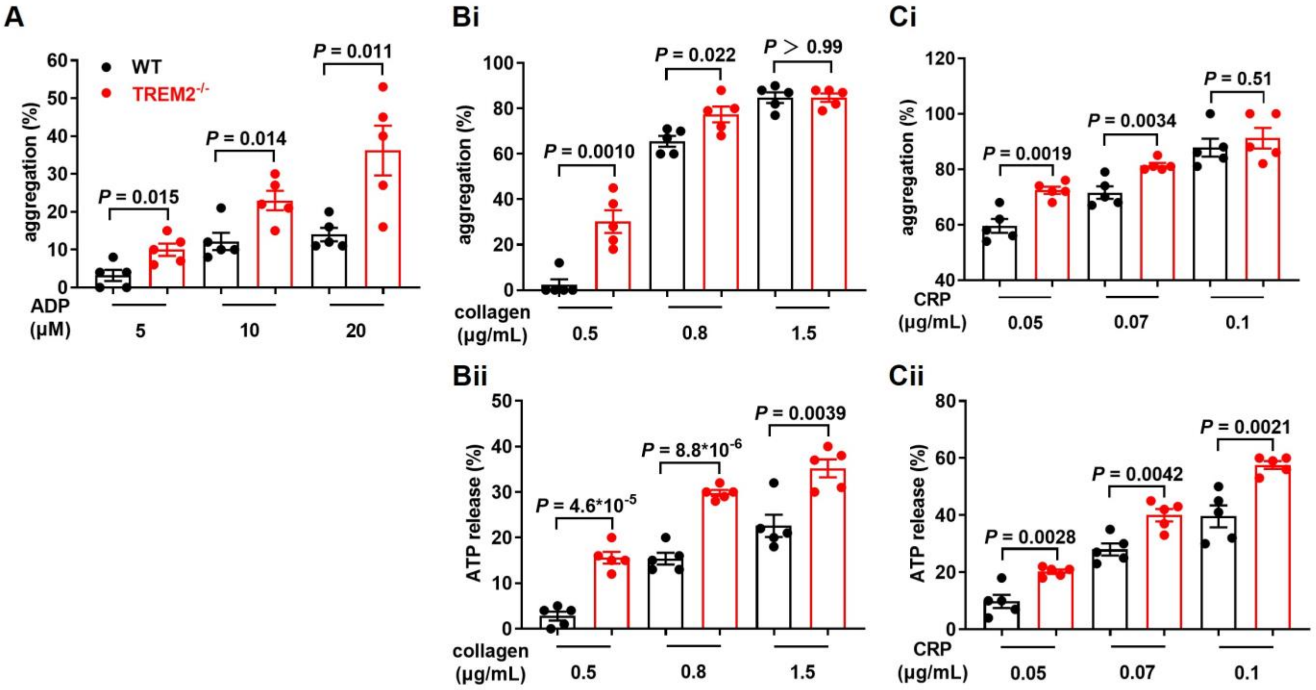


**Supplemental Figure 5. TREM2 deficiency enhances platelet aggregation and ATP release. A.** TREM2 deficiency increases platelet aggregation in response to ADP. Summary data of 5 experiments, corresponding to Figure 4A, are presented. **B.** TREM2 deficiency increases platelet aggregation (**Bi**) and ATP release (**Bii**) in response to collagen. Summary data of 5 experiments, corresponding to Figure 4A, are presented. **C.** TREM2 deficiency increases platelet aggregation (**Ci**) and ATP release (**Cii**) in response to CRP (n = 5). Summary data of 5 experiments, corresponding to Figure 4A, are presented. Unpaired t-test was performed in panel A, B, and C.

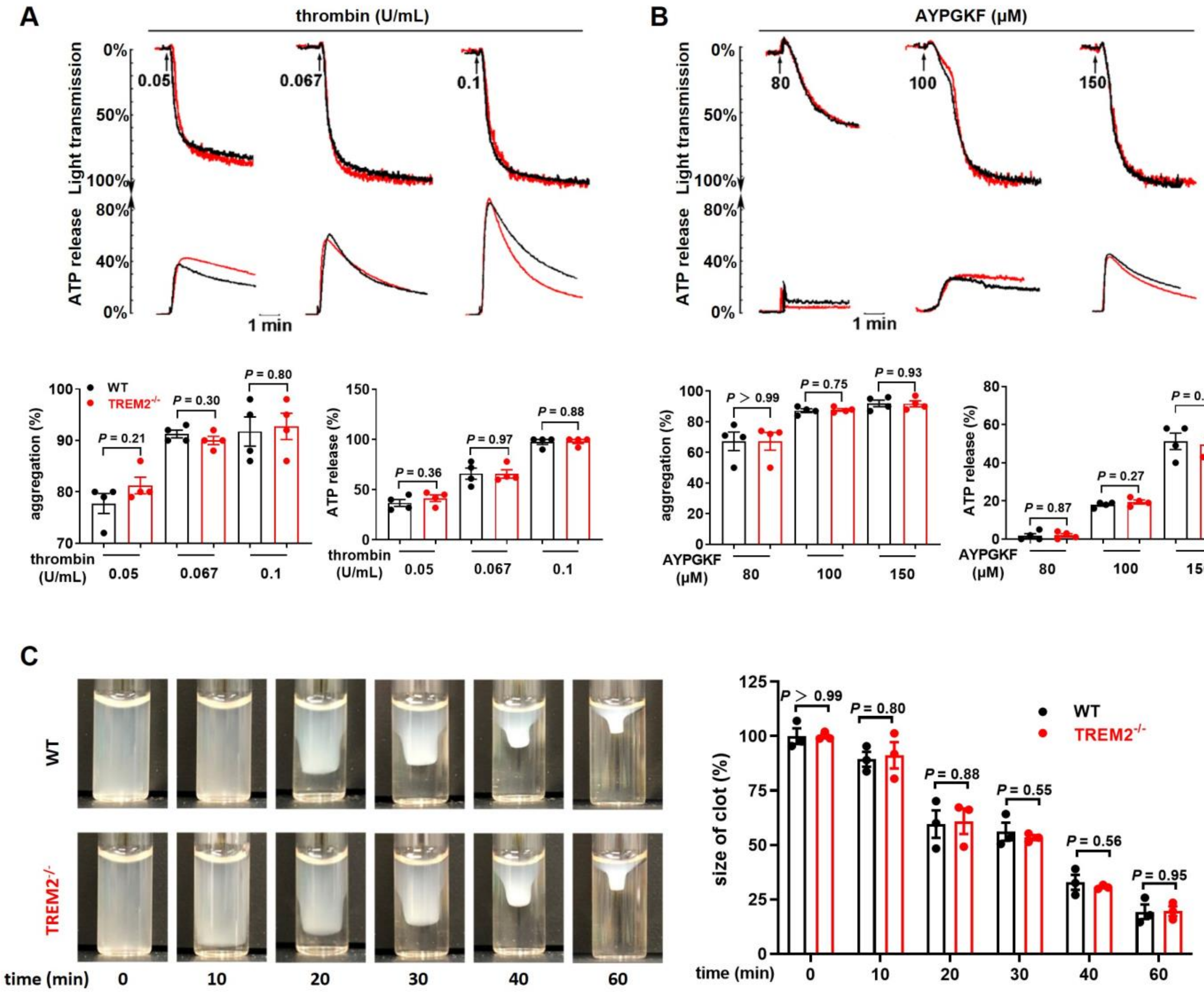


**Supplemental Figure 6. TREM2 deficiency does not affect either platelet aggregation and ATP release induced by thrombin and AYPGKF or clot retraction. A & B.** Washed platelets from wild-type (WT) and TREM2$^{-/-}$ mice were stimulated with thrombin (0.05, 0.067 or 0.1 U/mL), or AYPGKF (80, 100 or 150 μM). Aggregation and ATP release (with luciferase) were assessed under stirring at 1200 rpm. Typical tracings and the summary were provided (n = 4). **C.** Size of clots at indicated time was analyzed with ImageJ and the summary is shown (n = 3). Unpaired t-test was performed in panel A, B, and C.

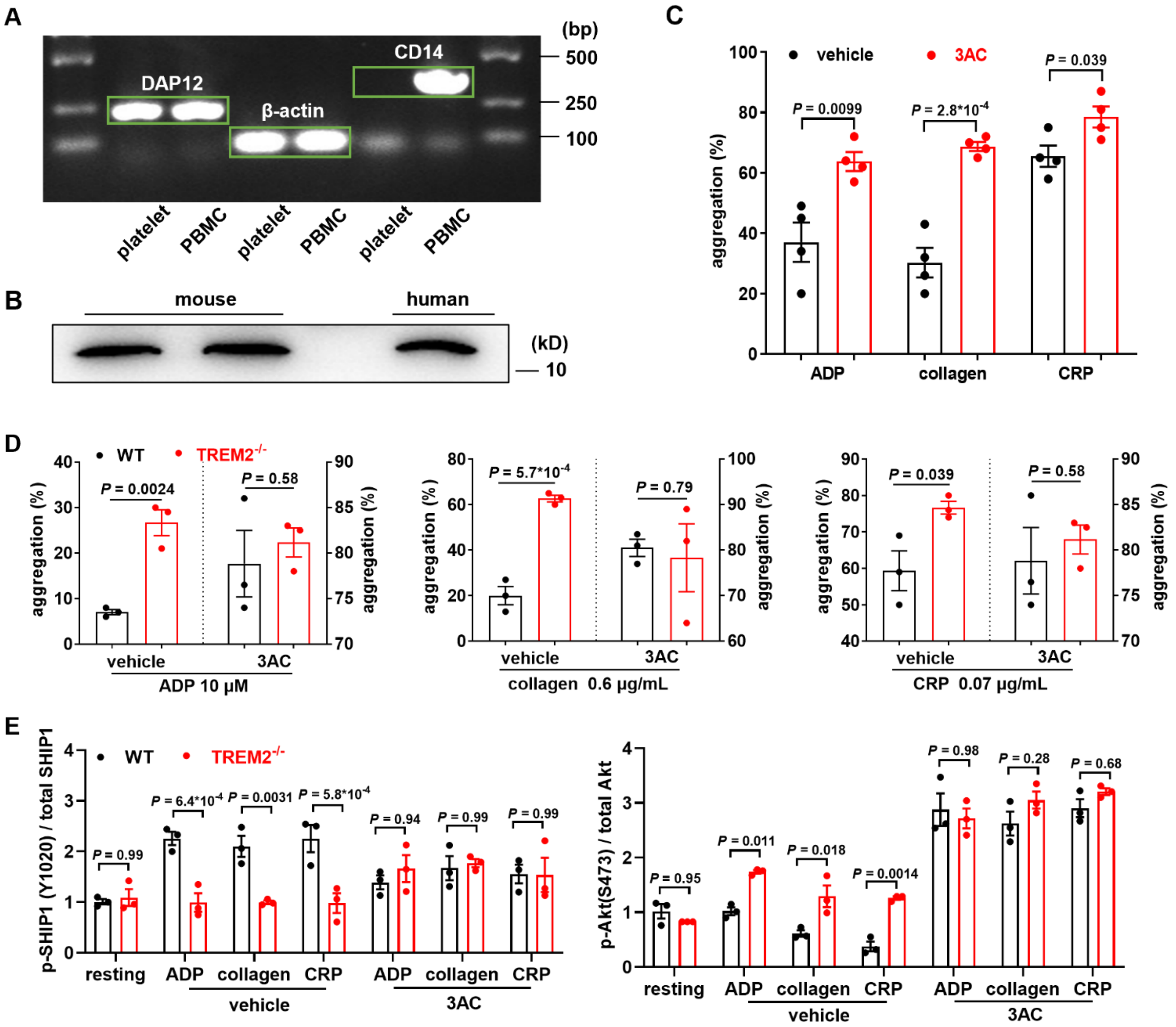


**Supplemental Figure 7. Platelets express DAP12, and TREM2 negatively regulates platelet activation by phosphorylating SHIP-1.** A. Human platelets express DAP12 detected by RT-PCR. Shown are 199 bp of DAP12, 357 bp of CD14 and 90 bp of β-actin. B. Platelets express DAP12 detected by Western blot. C. SHIP1 inhibitor 3AC (10 μM) enhances the aggregation of human washed platelets (n = 4). D. SHIP1 inhibitor 3AC potentiates WT and TREM2$^{-/-}$ platelet aggregation to the similar extent. Mouse washed platelets and 3AC 10 μM were used (n = 3). E. Similar phosphorylation levels of SHIP1 (Y1020) and Akt (S473) in activated WT and TREM2$^{-/-}$ platelets in the presence of SHIP1 inhibitor 3AC. Relative densities from 3 experiments, corresponding to Figure 7G, are presented. Two-way ANOVA followed by Sidak's multiple comparisons test was performed in panel E.

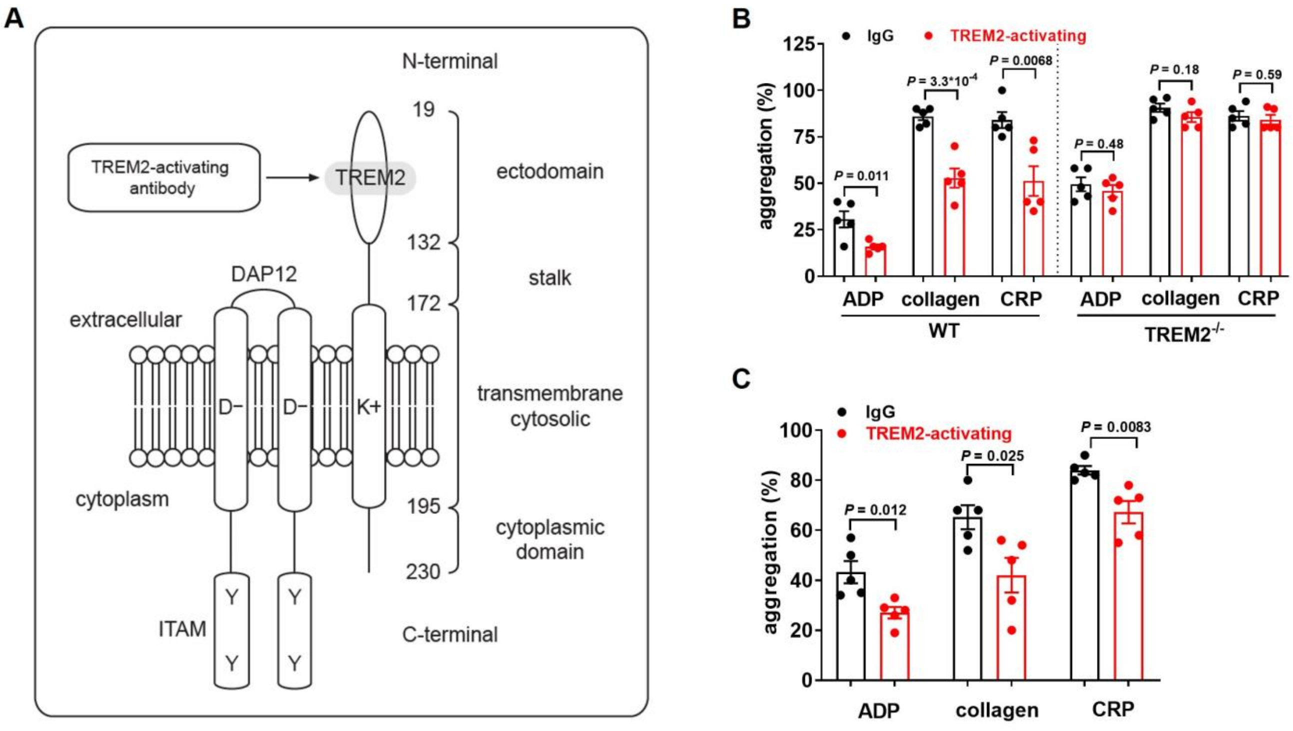


**Supplemental Figure 8. TREM2-activating antibody activates downstream signals of TREM2 and inhibits platelet aggregation TREM2-dependently. A.** Diagram showing the binding site of TREM2-activating antibody on TREM2. **B.** TREM2-activating antibody inhibits platelet aggregation of WT mice, but not TREM2$^{-/-}$ mice (n = 5). **C.** TREM2-activating antibody inhibits human platelet aggregation (n = 5). Unpaired t-test was performed in panel B and C.
