## Supplemental Tables for "ER stress-induced TREM2 downregulation exacerbates platelet activation and myocardial infarction in patients with coronary artery disease"

**Supplemental Table I. Primers used.**

| name | 5'-3' | experiment | species |
|---|---|---|---|
| β-actin-F | ggagattactgccctggctccta | qPCR | human |
| β-actin-R | gactcatcgtactcctgcttgctg | qPCR | human |
| TREM2-F | tcctgttgctggtcacagag | qPCR | human |
| TREM2-R | ctcccattctgcttcctcag | qPCR | human |
| GRP78-F | ccccgagaacacggtcttt | qPCR | human |
| GRP78-R | caaccaccttgaacggcaa | qPCR | human |
| CHOP-F | gcaagaggtcctgtcttcagatg | qPCR | human |
| CHOP-R | aagcagggtcaagagtggtga | qPCR | human |
| ATF4-F | ttctccagcgacaaggctaagg | qPCR | human |
| ATF4-R | ctccaacatccaatctgtcccg | qPCR | human |
| ATF6-F | cagacagtaccaacgcttatgcc | qPCR | human |
| ATF6-F | gcagaactccaggtgcttgaag | qPCR | human |
| Hsp60-F | tgccaatgctcaccgtaagcct | qPCR | human |
| Hsp60-R | agccttgactgccacaacctga | qPCR | human |
| HSPA4-F | gacctgccaatcgagaatcagc | qPCR | human |
| HSPA4-R | ctgcgttcttagcatcattccgc | qPCR | human |
| 1-1000-F | acggtaccggtccctgtgcttgga | plasmid construction | human |
| 1-1000-R | gtaagcttgccacccttccccagccaag | plasmid construction | human |
| 1001-2000-F | acggtacctttatagacttaaacatacac | plasmid construction | human |
| 1001-2000-R | gtaagcttcacgtgtgcaaaggc | plasmid construction | human |
| 2001-3000-F | gtggtaccattttcaacaaatac | plasmid construction | human |
| 2001-3000-R | taaagcttagctgccaaagt | plasmid construction | human |
| 1-200-F | acggtaccgcagtggccgactcctcctc | plasmid construction | human |
| 1-200-R | taaagcttgccacccttccccagcca | plasmid construction | human |
| 201-400-F | aaggtaccagatggcgggcat | plasmid construction | human |
| 201-400-R | taaagcttagccctggca | plasmid construction | human |
| 401-600-F | taggtaccagggaacagacacaggtagg | plasmid construction | human |
| 401-600-R | taaagctttctggtttcagccttgcaag | plasmid construction | human |
| 601-800-F | tgggtacccagaaaattccaggt | plasmid construction | human |

| | | | |
|---|---|---|---|
| 601-800-F | cgaagcttatctatacagt | plasmid construction | human |
| 801-1000-F | acggtaccggtccctgtgcttgga | plasmid construction | human |
| 801-1000-R | cgaagcttcccaagtttag | plasmid construction | human |
| TREM2-F | ccgctcgagtcccagcacaactccatttcccact | plasmid construction | human |
| TREM2-R | cccaagcttgtcaggcgctgcccacaggaact | plasmid construction | human |
| TREM2-mutation-F | atggcgggcagctggtggagggtctgaatacagctgtgaggatag | plasmid construction | human |
| TREM2-mutation-R | ctccaccagctgcccgccatcttctggtttcagccttgcaaga | plasmid construction | human |
| sh-chop-F | gatccggtcctgtcttcagatgaattcaagagattcatctgaagacaggaccttttttg | plasmid construction | human |
| sh-chop-R | aattcaaaaaaggtcctgtcttcagatgaatctcttgaattcatctgaagacaggaccg | plasmid construction | human |
| TREM2-F | tttccttgcagagcctag | ChIP | human |
| TREM2-R | aagaggtgctggatgagg | ChIP | human |
| GAPDH-F | tactagcggttttacgggcg | ChIP | human |
| GAPDH-R | tcgaacaggaggagcagagagcga | ChIP | human |
| GAPDH-F | tgaccagaacatcatccctg | qPCR | mouse |
| GAPDH-R | tcagatccacgacggacaca | qPCR | mouse |
| TREM2-F | agtcatcgagtttcgagggtg | qPCR | mouse |
| TREM2-R | aaggtggtaggctagaggtga | qPCR | mouse |
| TREM2-mutant R | caataagacctggcacaagga | Genotyping | mouse |
| TREM2-common F | tcagggagtcagtcattaacca | Genotyping | mouse |
| TREM2-wild-type R | agtgcttcaaggcgtcataagt | Genotyping | mouse |
| β-actin-F | gtcattccaaatatgagatgcgttg | RT-PCR | human |
| β-actin-R | ggactgggccattctccttag | RT-PCR | human |
| TREM2-F | acgagatcttgcacaaggca | RT-PCR | human |
| TREM2-R | gatggaagtgggtgggaagg | RT-PCR | human |
| CD14-F | cagtagcttgacacggtcaag | RT-PCR | human |
| CD14-R | tccaggattgtcagacaggtc. | RT-PCR | human |
| β-actin-F | acccataccgtcgtccaatc | RT-PCR | mouse |
| β-actin-R | ccccttcaggttgcaaatcg | RT-PCR | mouse |
| TREM2-F | acagcacctccaggaatcaag | RT-PCR | mouse |
| TREM2-R | cctggctggacttaagctgt | RT-PCR | mouse |

| | | | |
|---|---|---|---|
| CD14-F | cccagtcagctaaactcgct | RT-PCR | mouse |
| CD14-R | ccactgcttgggatgatgga | RT-PCR | mouse |

**Supplemental Table II. Antibodies used.**

| Name | assay | cat# | company |
|---|---|---|---|
| CD41[EPR4330] | IF | ab134131 | Abcam |
| TREM2 (D8I4C) | IF | #91068 | CST |
| DAPI | IF | C1002 | Beyotime |
| IL-1 beta | IF | ab254195 | Abcam |
| TNF-α (D2D4) XP | IF | #11948 | CST |
| TREM2- N-terminal | WB | ab175262 | Abcam |
| GAPDH | WB | ab181602 | Abcam |
| CHOP (L63F7) | WB/IF | #2895 | CST |
| GRP78 bip | WB/IF | ab21685 | Abcam |
| TREM2 | IP | ab209814 | Abcam |
| DAP12 (D7G1X) | WB | #12492 | CST |
| p-SHIP1(Y1020) | WB | #3941 | CST |
| Akt (pan) (11E7) | WB | #4685 | CST |
| p-Akt (S473) (D9E) | WB | #4060 | CST |
| SHIP1 | WB | SC-8425 | SC |
| C/EBPα | WB | #8178 | CST |
| p-β3 (Y747) | WB | 11060 | SAB |
| p-β3 (Y759) | WB | 11282 | SAB |
| β3 | WB | 18309-AP-1 | Proteintech |
| p-Src (Y416) | WB | 2101S | CST |
| Src | WB | A19119 | Abclonal |
| p-P38 (Y182) | WB | sc-166182 | Santa |
| P38 | WB | 14064-1-AP | Proteintech |
| TREM2[6E9] (FITC) | flow cytometry | ab236286 | Abcam |
| CD62P | flow cytometry | 561923 | BD |
| JoN/A | flow cytometry | #D200 | Emfret |
| DDIT3 (CHOP) | EMSA | ab117195 | Abcam |
| C/EBPα | EMSA | ab51301 | Abcam |
| rabbit anti-mouse thrombocyte | eliminate platelet | J1943 | Accurate Chemical |
| actin-tracker green | spreading | C1033 | Beyotime |

| | | | |
|---|---|---|---|
| C/EBPα | CHIP/EMSA | SC-365318 | SC |
| DAP12 | WB/IP | #12492 | CST |
| p-Tyr-100 | WB | #9411 | CST |
| TREM2 (E6T1P) | TREM2-activating antibody | #61788 | CST |
| Rabbit IgG (DA1E) | isotype control | #3900 | CST |
| Cd42b | IHC | 12860-1-AP | Proteintech |

**Supplemental Table III. Characteristics of study population in Figure 1D, 2D, and 2G.**

| | healthy | SAP | ACS | *P* value |
|---|---|---|---|---|
| n | 19 | 12 | 16 | |
| male | 16 (84.2%) | 10 (83.3%) | 14 (87.5%) | |
| female | 3 (15.8%) | 2 (16.7%) | 2 (12.5%) | 0.9445 |
| age | 68.6 ± 15.4 | 63.0 ± 7.8 | 65.5 ± 9.9 | 0.4468 |
| hypertension | NA | 7 (58.3%) | 9 (56.3%) | > 0.99 |
| smoking | NA | 4 (33.3%) | 4 (25.0%) | 0.6908 |
| total cholesterol (mmol/L) | NA | 3.752 ± 1.729 | 3.998 ± 1.630 | 0.7035 |
| triglyceride (mmol/L) | NA | 3.531 ± 4.907 | 2.133 ± 2.682 | 0.3425 |
| platelet count (× $10^9$ /L) | NA | 192.4 ± 55.4 | 172.9 ± 48.2 | 0.3296 |
| hs-CRP (mg/L) | NA | 23.34 ± 46.55 | 33.93 ± 72.97 | 0.6622 |

SAP: stable angina pectoris; ACS: acute coronary syndrome; NA: not applicable. Data were presented as mean ± SEM. The difference of gender was analyzed by Chi-square test and the difference of age was analyzed by One-way ANOVA followed by Kruskal-Wallis test. Differences in other data were analyzed using unpaired t-test.

**Supplemental Table IV. Characteristics of study population in Figure 1E.**

| | healthy | SAP | ACS | *P* value |
|---|---|---|---|---|
| n | 32 | 31 | 65 | |
| male | 21 (65.6%) | 21 (67.8%) | 49 (75.4%) | |
| female | 11 (34.4%) | 10 (32.2%) | 16 (24.6%) | 0.5440 |
| age | 63.2 ± 13.7 | 66.9 ± 12.1 | 65.9 ± 14.6 | 0.5396 |
| hypertension | NA | 19 (61.3%) | 36 (55.4%) | 0.6629 |
| smoking | NA | 12 (38.7%) | 21 (32.3%) | >0.99 |
| total cholesterol (mmol/L) | NA | 3.166 ± 1.527 | 3.580 ± 1.710 | 0.2546 |
| triglyceride (mmol/L) | NA | 1.677 ± 1.273 | 1.551 ± 0.952 | 0.5876 |
| platelet count (× $10^9$ /L) | NA | 190.1 ± 52.4 | 202.7 ± 69.1 | 0.3688 |
| hs-CRP (mg/L) | NA | 4.574 ± 4.872 | 32.74 ± 47.71 | 0.0015 |

SAP: stable angina pectoris; ACS: acute coronary syndrome; NA: not applicable. Data were presented as mean ± SEM. The difference of gender was analyzed by Chi-square test and the difference of age was analyzed by One-way ANOVA followed by unpaired test. Differences in other data were analyzed using unpaired t-test.

**Supplemental Table V. Characteristics of the study population in Figure 1F.**

| | healthy | SAP | ACS | *P* value |
|---|---|---|---|---|
| n | 25 | 25 | 25 | |
| male | 16 (40.0%) | 17 (68.0%) | 17 (68.0%) | |
| female | 9 (60.0%) | 8 (34.0%) | 8 (34.0%) | 0.9418 |
| age | 62.7 ± 1.8 | 63.6± 1.9 | 66.3 ± 1.8 | 0.4491 |

SAP: stable angina pectoris; ACS: acute coronary syndrome. Data were presented as mean ± SEM. The difference of gender was analyzed by Chi-square test and the difference of age was analyzed by One-way ANOVA followed by unpaired test.

**Supplemental Table VI. Characteristics of study population in Supplemental Figure 1.**

| n | 50 |
|---|---|
| age | 67.28 ± 12.92 |
| male (%) | 33 (66.00%) |
| hypertension (%) | 30 (60.00%) |
| smoking (%) | 13 (26.00%) |
| total cholesterol (mmol/L) | 4.331 ± 1.511 |
| triglyceride (mmol/L) | 2.121 ± 2.294 |
| platelet count (× $10^9$ /L) | 201.9 ± 64.88 |
| hs-CRP (mg/L) | 38.18 ± 57.77 |

Data were presented as mean ± SEM.

**Supplemental Table VII. Hematologic parameters of TREM2 deficient mice.**

| | WT (n = 26) | TREM2$^{-/-}$ (n = 23) | *P* value |
|---|---|---|---|
| WBC (× $10^9$ /L) | 2.80 ± 0.23 | 2.23 ± 0.22 | 0.0562 |
| RBC (× $10^{12}$ /L) | 7.56 ± 0.16 | 7.71 ± 0.22 | 0.4825 |
| PLT (× $10^{11}$ /L) | 6.27 ± 0.24 | 8.07 ± 0.55 | 0.0055 |
| MPV (fL) | 3.80 ± 0.23 | 4.04 ± 0.10 | 0.0356 |
| PDW (fL) | 16.20 ± 0.39 | 16.40 ± 0.09 | 0.2572 |

WBC: white blood cell; RBC: red blood cell; PLT: platelet; MPV: mean platelet volume; PDW: platelet distribution width. Data were presented as mean ± SEM and analyzed by unpaired t-test.
